# Horizontal transmission of functionally diverse transposons is a major source of new introns

**DOI:** 10.1101/2024.06.04.597373

**Authors:** Landen Gozashti, Anne Nakamoto, Shelbi Russell, Russ Corbett-Detig

## Abstract

Specialized transposable elements (TEs), introners, are one of the major drivers of intron gain in diverse eukaryotes. However, the molecular mechanism(s) and evolutionary processes driving introner propagation within and between lineages remain elusive. We analyze 8716 genomes, revealing 1093 introner families in 201 species spanning 1.7 billion years of evolution. Introners are derived from functionally diverse TEs including families of terminal-inverted-repeat DNA TEs, retrotransposons, cryptons, and helitrons as well as mobile elements with unknown molecular mechanisms. We identify eight cases where introners recently transferred between divergent host species, and show that giant viruses that integrate into genomes may facilitate introner transfer across lineages. We propose that intron gain is an inevitable consequence of TE activity in eukaryotic lineages, thereby resolving a key mystery of genome structure evolution.

## Introduction

Spliceosomal introns are a fundamental component of eukaryotic genomes with poorly understood origins. Introns are noncoding regions within genes that must be removed from transcripts before translation. They are nearly ubiquitous across eukaryotic genomes and contribute to a range of molecular processes including regulation of transcription and protection against transcription-associated genome instability (*1*, *2*). Introns also enable alternative splicing, vastly expanding the molecular and functional diversity encoded by eukaryotic genes (*3*). Many introns are essential to gene function and intron deletion can be costly and even lethal (*4*). Although introns now serve important molecular functions, these roles must have evolved after the emergence of introns (*5*). Therefore, the proximal origins of spliceosomal introns remain poorly understood, representing a longstanding question in biology.

Transposable elements (TEs) – selfish genes that propage within genomes by copying themselves – may be the source of most new introns. The number of introns per gene varies tremendously across species (range ∼0.003-20 per gene) (*6*–*8*), and comparisons of orthologous intron positions across lineages suggest a history of rapid episodic intron gain (*7*). Several mechanisms of intron gain have been proposed (reviewed in (*9*)); however, *de novo* intron creation by specialized intron-generating transposable elements, termed introners, is the only mechanism that could explain the “bursts” of intron gains observed across lineages (*7*).

Introners are transposable elements which can be correctly spliced out of exons upon insertion (either by encoding or co-opting spliceosomal recognition sequences), and have the unique ability to create thousands of introns within a single genome (*10*, *11*). Recently-active introners can be identified by comparing sequence similarity among introns from diverse locations across the genome, have been reported in diverse lineages and may explain the vast majority of ongoing intron gain (*12*–*18*). However, the mechanisms driving introner proliferation remain almost entirely unexplored, and it is unclear whether introners arise from diverse transposable elements or are restricted to a specific mechanism of mobilization. Furthermore, introners show patchy taxonomic distributions and are enriched in species that undergo frequent horizontal gene transfer (HGT), such as aquatic unicellulars and fungi, suggesting that HGT may play an important role in shaping introner distributions across the tree of life (*12*). Nonetheless, direct evidence for HGT has not been discovered and the precise molecular mechanisms of transposition are almost completely unknown. Thus, the biological processes underlying *de novo* intron creation remain poorly understood.

### Transposable elements generate introns across the eukaryotic tree

We systematically searched for introners in 8716 annotated eukaryotic genome assemblies (see Methods) and discovered an unprecedented diversity of species whose genomes contain introns derived from recent transposition. Introners are present in an exceptionally broad range of eukaryotic species (Fig. 1; table S1). In line with prior work, we discovered an abundance of introners in aquatic organisms, unicellular species and fungi (>98% of introner-containing species are aquatic, unicellular or fungi; table S1) (*12*). However, we also find introner families in an expanded range of taxa (see Methods; fig. S1-S7). In particular, we observe recently-active introners in land plants including the grasses, (*Panicum virgatum*) and eudicots (*e*.*g*., *Salvia splendens*), as well as an echinoderm, the purple sea urchin (*Strongylocentrotus purpuratus*). Our search also revealed introners in basidiomycete fungi as well as a myriad of diverse protist lineages (Fig. 1; table S1). Thus introner-derived introns are widespread across several previously unexplored taxa, highlighting the universality of this mechanism of intron gain.

**Fig 1.**
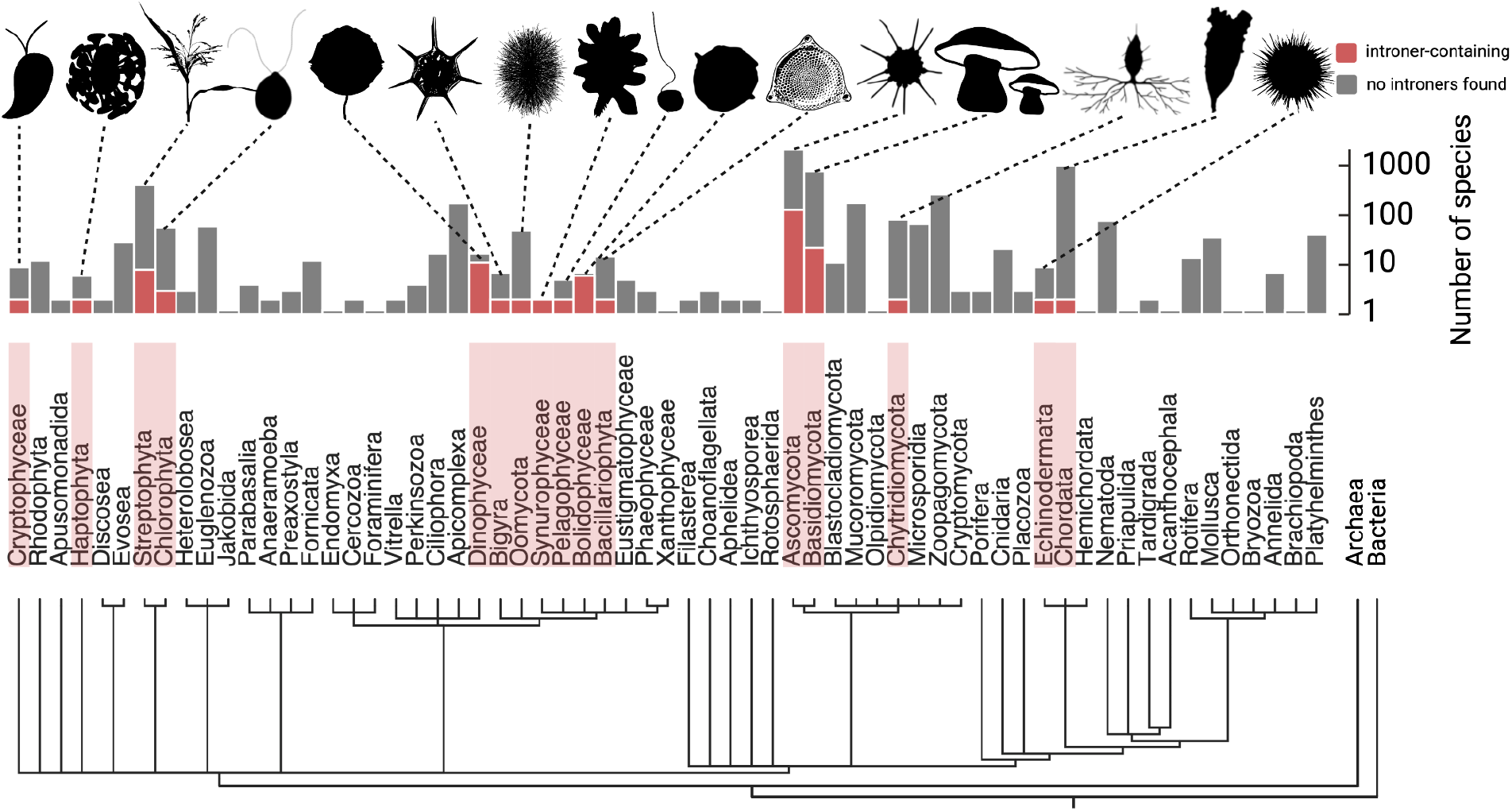
Distribution of Introners across the eukaryotic tree of life. Relationships between phyla for considered genomes are shown as a cladogram. Considered genomes are arranged into phyla according to NCBI taxonomy with outgroup lineages that do not contain a spliceosome: archaea and bacteria. Phyla with introner-containing species are highlighted in red. Gray and red bars display the number of species evaluated and the number of introner-containing species we discovered for each phylum, respectively. Silhouettes show examples of introner-containing species for each phylum and were retrieved from PhyloPic.

### Intron-generating TEs span exceptional mechanistic diversity

Mirroring the broad host taxonomic variety, TEs capable of generating introns are also exceptionally mechanistically and evolutionarily diverse. We used a combination of systematic and manual approaches to identify transposition mechanisms and classify newly identified introners as they relate to known transposable elements. Our approach classified TEs using homology to known TEs, expected functional domains and structural features consistent with specific mechanisms, and conserved terminal motifs observed for many elements (See Methods; table S2). We find that introners arise from TEs spanning ∼90% of orders and at least 50% of superfamilies of known mobile genetic elements as defined by (*19*) (Fig. 2A). These include diverse TIR DNA transposons, LTR retrotransposons, nonLTR retrotransposons, rolling circle helitrons and tyrosine recombinase (Crypton) elements (each comprising ∼79%, ∼14%, 5%, 2% and <1% of confidently categorized examples; Fig. 2B; table S2; reviewed in (*20*, *21*)).

**Fig 2.**
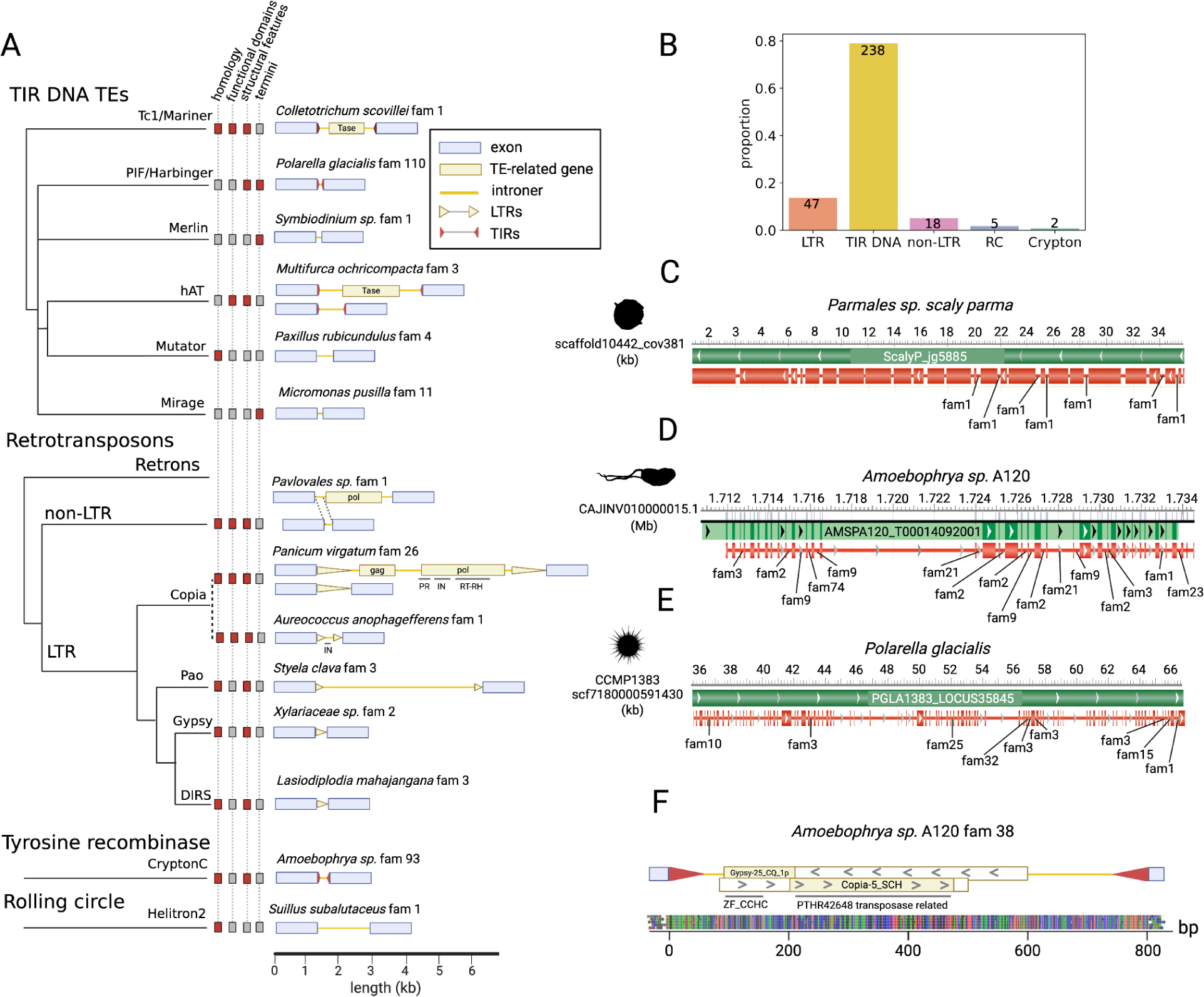
Functional diversity and exemplary introners for major TE families. **(A)** Cladogram displaying relationships among introner transposable elements families (following (*20*)), and examples of the introner structure for each family discovered in this work. Cladogram leaf tips denote lines of evidence supporting introner classifications, including homology to known TEs, expected functional domains, structural features such as LTRs or TIRs and conserved termini (see Methods; Tase = Transposase). For introner structures, blue boxes denote flanking host exons. Introner features are shown in gold, with the exception of TIRs, which are plotted in red for visibility. **(B)** Proportion of introner families represented by each major TE subclass for subset of introners which could be confidently classified. Numbers on bars display total counts for each subclass. **(C-E)** Genome browser screen-shots for selected introner-containing genes, generated using NCBI’s Genome Data Viewer. Introner positions are annotated for different introner families in each species (table S3). Species silhouettes were retrieved from PhyloPic. **(F)** Model of *Amoebophrya* sp. A120 family 38. Introner ORFs are shown as boxes with regions showing homology to known TEs shaded in gold. Functional domains spanning these regions of homology are labeled below, followed by a multiple sequence alignment of all intron-generating copies from this introner family.

Remarkably, many of these transposable elements do not share an inferable common ancestor, suggesting origins that predate eukaryotes themselves (*20*, *22*, *23*). Additionally, ∼72% of elements cannot be confidently categorized. We suspect that these unknown elements will provide insight into as yet unexplored mobile elements (table S2). This breadth emphasizes the unprecedented functional diversity of transposons that are capable of generating introns on genomic scales.

We further identified elements where both autonomous and non-autonomous elements contribute concurrently to intron gain (Fig. 2A; table S2). For example, an introner family in *Panicum virgatum* displays homology to known Copia LTR elements and several copies display functional domains required for autonomous Copia mobilization. However, other copies exist as solo-LTRs, products of ectopic recombination resulting in the removal of the internal TE sequence (*24*) (Fig. 2A). Both fully-autonomous elements and solo-LTRs generate functional introns. TIR DNA transposons display similar patterns in several species (Fig. 2A).

In some species a single introner family contributes to the vast majority of introner-mediated intron gain, whereas other species display multiple abundant introners with divergent origins. For example, in the marine diatom *Parmales sp. scaly parma*, one introner family contributes to >91% of recognizable intron gains via an unknown mechanism (Fig. 2C). In contrast, the parasitic dinoflagellate *Amoebophrya* sp. A120 harbors introners attributed to nonLTR retrotransposons, LTR retrotransposons, TIR DNA transposons and Cryptons (table S2). These diverse introners contribute to massive intron gain in *Amoebophrya*, in some cases generating tens of introns in a single gene (Fig. 2D). Similarly to *Amoebophrya*, the free living dinoflagellate, *Polarella glacialis*, harbors a tremendous diversity of introners contributing to ongoing intron gain (Fig. 2E; table S2).

Beyond transposon families that we can confidently identify, some introners are derived from previously unknown transposition mechanisms. In particular, in *Amoebophrya* sp. A120, one intron-generating transposon family displays hallmarks of DNA transposons but shares homology to LTR retrotransposons transposons. This element contains open reading frames with homology to *Copia* and *Gypsy* LTR elements, but is flanked by DNA element-like terminal inverted repeats and exhibits several near-identical copies (Fig. 2F; fig. S8). Interestingly, this homologous region within *Copia-5_SCH* spans part of a putative transposase as annotated by Panther (PTHR42648), but does not overlap with an integrase or any other specific functional domain associated with DNA transposons (fig. S9). This unexpected discovery might provide clues to the ancient relationships among LTR retrotransposons and DNA transposons (*25*, *26*), and further highlights the still-unknown diversity of transposable elements that are apparently capable of generating new introns across genomes.

### Horizontal gene transfer shapes introner distributions

Evidence of recent homology strongly implicates horizontal transmission as one of the major drivers of intron propagation within and among eukaryotic lineages. Transposable elements frequently horizontally transfer between lineages (*27*, *28*), suggesting that this may be an important phenomenon shaping introner evolution. By performing an extensive blast search of all introner families discovered in this work against the NCBI nucleotide database (see Methods; table S4), we found 8 unambiguous examples of recent horizontal transmission of introner generating transposons (4% of all introner-containing species; table S5). Horizontal transmission of transposable elements is therefore a critical biological process driving intron propagation between divergent populations.

We identified two examples of HGT within ascomycete fungi where the same elements have recently generated introns among highly divergent lineages (Fig. 3A). *Xylaria* and *Lasiodiplodia* last shared a common ancestor approximately 350 MYA (*29*, *30*). Nonetheless, each species’ genome contains an introner family with 83.8% sequence similarity across the complete 134bp sequence between elements found in either lineage (Fig. 3B; table S4). Furthermore, molecular dating estimates suggest that introners in these distant lineages last diverged a mere ∼7 MYA (Fig. 3B; fig S10). Similarly, *Alternaria* and *Parastagonospora* last shared a common ancestor approximately 134 MYA (Fig. 3A), but each contains a recently active introner family of length 64bp with ∼78.5% sequence similarity (table S4) (*29*, *31*). Estimates for *Alternaria* and *Parastagonospora* introners suggest they last diverged ∼10 MYA (Fig. 3C; fig. S11). Notably, we identified introners in multiple genomes from different strains or species within these two genera, and introners found in each genome are most closely related to others found in genomes of the same genus. This implies that introner observations are not an idiosyncratic assembly artifact and strongly implicates recent transposition in highly divergent lineages rather than the transfer of an intron that did not continue transposing in the recipient population (Fig. 3C).

**Fig. 3:**
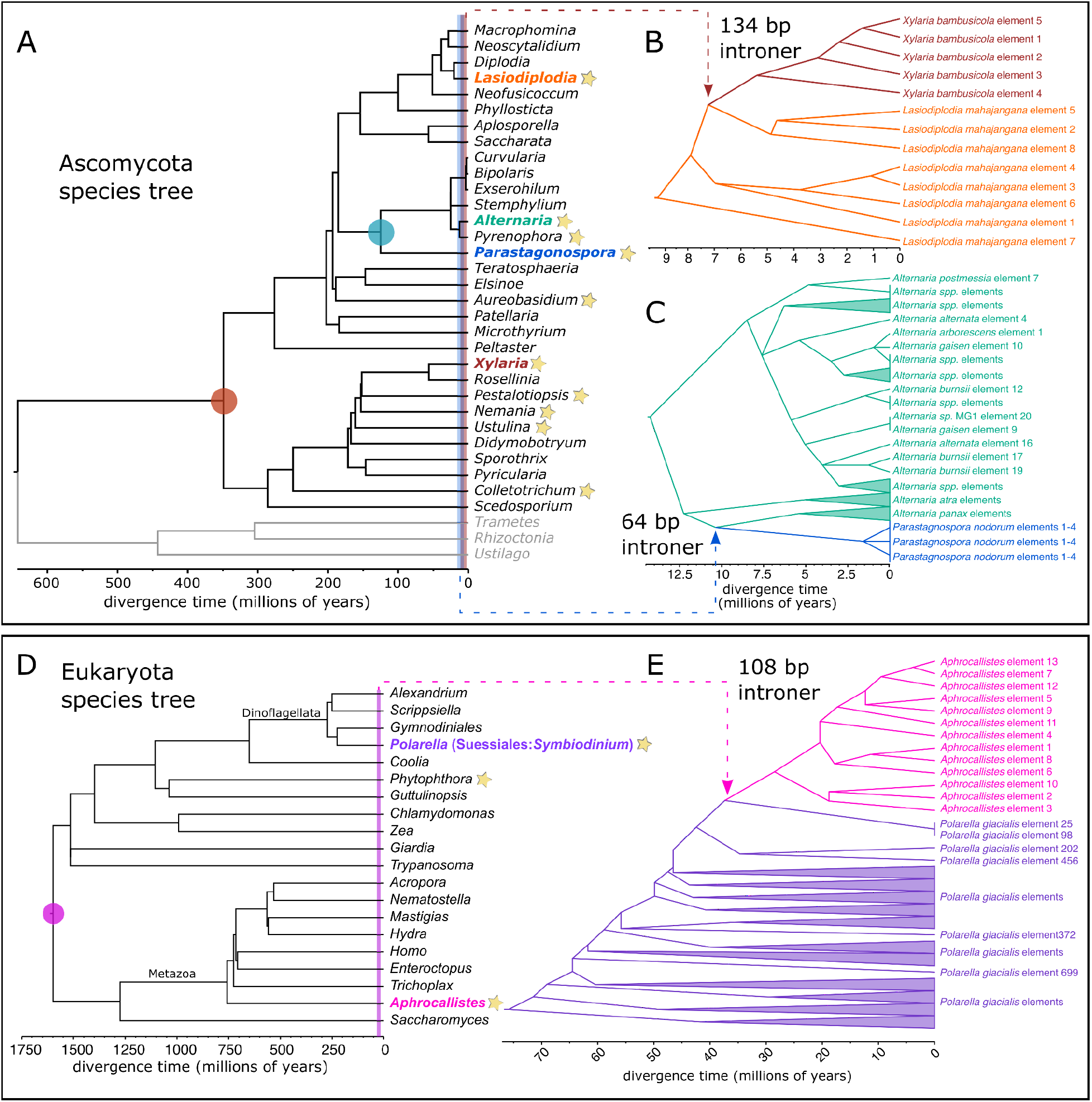
Horizontal gene transfer of introners between divergent taxa. **(A)** Phylogeny of ascomycete fungi retrieved from TimeTree (*29*). Lineages showing evidence of HGT for introners are highlighted with colored font. Shaded circles denote the last common ancestor for species showing introner HGT and vertical lines show estimated divergence times for homologous introners between lineages with brown and blue corresponding to *Xylaria*+*Lasiodiplodia* and *Parastagonospora*+*Alternaria*, respectively. Stars indicate other genera where we identified introners (table S6). **(B-C)** Phylogenies for prospective horizontally transferred introner families in (**B)** *Xylaria* and *Lasiodiplodia*, and (**C)** *Parastagonospora* and *Alternaria*. Colors in (**B**) and (**C**) correspond to leaf labels in (**A**). (**D**) Phylogeny of eukaryotes retrieved from TimeTree (table S7) (*29*). A shaded circle denotes the last common ancestor between the introner-containing dinoflagellate lineage *Polarella* (purple) and the sponge lineage *Aphrocallistes* (pink), which contains homologous sequences but no evidence of introner-mediated intron gain. A shaded line denotes the estimated divergence between *Polarella* introners and *Aphrocallistes* homologs. Stars indicate other genera where we identified introners (table S7). (**E**) Phylogeny of *Polarella* introners and *Aphrocallistes* homologs.

We also discovered HGT between lineages in which introners in one species’ genome may propagate as ordinary transposable elements in another. Introners in the dinoflagellate *Polarella glacialis* show strong homology to multiple loci in the glass sponge *Aphrocallistes beatrix* (up to 72% sequence similarity and 87% coverage; tables S4-S5, S8-S9; fig. S12). However, we find no evidence that these introner homologs contribute to intron gain in *A. beatrix*. The common ancestor between dinoflagellates and sponges is also the last eukaryotic common ancestor (>1.6 BYA (*29*, *32*)) (Fig. 3D). Nonetheless, estimates suggest that *P. glacialis* introners and their *A. beatrix* homologs diverged less than one million years ago (Fig. 3E). *P. glacialis* and *A. beatrix* share overlapping ranges in both arctic and antarctic regions (*33*–*35*). Furthermore, dinoflagellates and sponges exhibit frequent horizontal gene transfer, including inter-kingdom and inter-domain transfer (*36*–*38*) (supplementary text; fig. S14), implying that introner gain through HGT may be driven by these species’ ecology (*39*, *40*). These results imply that HGT of introners can occur between even the most divergent eukaryotic lineages.

### Giant viruses may facilitate horizontal gene transfer of introners across lineages

Viruses are frequent vectors of horizontal transfer and may underlie transmission of TEs between diverse taxa (*41*–*47*). A bioinformatic search across our dataset revealed several cases of introners within viral genes integrated within host genomes (table S10; see Methods). An introner-containing gene in *Alternaria burnsii* (GeneID:62205305) is homologous to a klosneuvirus protein (MK072332.1; ∼50% Sequence ID, e-value∼0), and the upstream protein also shares high levels of homology to a klosneuvirus virus (MK072078.1; e-value = 6.28E-17). Klosneuviruses are a clade of giant viruses that infect diverse eukaryotic hosts including fungi and protists, acquire host genes at high rates, and contribute to HGT (*47*, *48*). Other examples of introner-containing genes homologous to klosneuviruses include the divergent dinoflagellates *Effrenium voratum* (gene EVOC421_LOCUS2994 and viral accession MT418680.1, ∼54% SID, e-value∼0) and *Polarella glacialis* (gene PGLA1383_LOCUS8179 and viral accession KY684104.1.1, 91% SID, e-value=2.42E-117). Overall, introner-containing genes are enriched for viral proteins (tblastn e-values<1E-5; MWU, p<0.0001 compared to expectations from random resampling). Among these, klosneuviruses are strongly overrepresented (9.4% of introner-containing genes that are viral in origin, whereas klosneuviruses comprise <0.0001 of sequences in RVDB; binomial test p<0.000001). These findings suggest that viruses infecting diverse hosts may provide a mechanistic explanation for HGT of introners across divergent lineages.

## Discussion

Our comprehensive survey of the tree of life provides direct evidence that HGT of transposable elements contributes to intron gain across eukaryotic lineages. Although other factors must contribute, we propose that HGT explains several foundational patterns in intron evolution: (1) the genome of the last eukaryotic common ancestor, which was a single-celled aquatic organism, contained thousands of introns, possibly due to an abundance of HGT of diverse mobile genetic elements (*23*, *49*, *50*); (2) introners and intron gain are disproportionately abundant in the genomes of aquatic and fungal taxa – both are known to experience high rates of HGT (*12*, *27*, *40*, *51*, *52*); and (3) conversely, the apparent lack of recent intron gain in lineages such as vertebrates (*53*, *54*) may in part reflect the paucity of HGT and relative lack of TE diversity for many vertebrate lineages (*51*, *55*, *56*). Therefore widespread HGT of intron generating transposons resolves a fundamental question about why new introns evolve and what determines their abundances across species.

The abundance of evolutionarily and mechanistically diverse transposable elements that generate introns across the eukaryotic tree of life illuminates a longstanding mystery of evolution. Our results demonstrate that seemingly any variety of transposable element can and does generate introns within eukaryotic genomes. Efficient splicing after insertion into an exon should drastically reduce a TE’s deleterious effect on host fitness, and provides an adaptation both from the perspective of a transposable element and its host. Selective pressure for this innovation should be most pronounced when a TE is highly active, such as after the introduction of a “new” TE unrecognizable to host suppression machinery via HGT (*28*). Thus, HGT may not only facilitate introner transmission between divergent lineages but also favor their evolution *de novo* from diverse TEs. The fundamental implication is that introns will accumulate as an inevitable consequence of the ubiquitous genetic conflict between eukaryotic genomes and transposable elements.

## Acknowledgements

L.G. thanks Hopi E. Hoekstra for her support and advice throughout this work. The computations for this work were run on the FASRC Cannon cluster supported by the FAS Division of Science Research Computing Group at Harvard University. The authors thank Erik Enbody, Jennifer Chen, Peter Sudmant, and David Haussler for helpful feedback on this manuscript.

## Funding

This work was supported in part by R35GM128932 to R.C.-D. and R00GM135583 to S.L.R. A.N. was supported by an NSF GRFP.

## Authors contributions

L.G. and R.C.-D. conceived and designed the research. L.G. and A.N. performed introner discovery and classification. L.G., A.N. and S.L.R. analyzed cases of horizontal gene transfer between divergent lineages. L.G., A.N., and S.L.R. contributed to data visualization. L.G. and R.C.-D. provided resources for this work. R.C.-D. advised the research. All authors wrote the manuscript. All authors edited and contributed to the manuscript revision.

## Competing interests

The authors declare no competing interests.

## Data and materials availability

Our introner discovery pipeline is available at https://github.com/lgozasht/Introner-elements. Consensus sequences for all introner families are also available at https://github.com/lgozasht/Introner-elements.

## Supplementary Materials

Materials and Methods

Supplementary Text

Figs. S1 to S15

Tables S1 and S13

References (56-101)

## Notes

### Competing Interest Statement

The authors have declared no competing interest.

### Summary of Updates

We fixed a mistake in figure 2B where proportions of introners were displayed rather than proportions of introner families.

## References and notes

1. A. Bonnet, A. R. Grosso, A. Elkaoutari, E. Coleno, A. Presle, S. C. Sridhara, G. Janbon, V. Géli, S. F. de Almeida, B. Palancade, Introns protect eukaryotic genomes from transcription-associated genetic instability. Mol. Cell 67, 608–621.e6 (2017).

2. O. Shaul, How introns enhance gene expression. Int. J. Biochem. Cell Biol. 91, 145–155 (2017).

3. M. Irimia, S. W. Roy, Origin of spliceosomal introns and alternative splicing. Cold Spring Harb. Perspect. Biol. 6 (2014).

4. J. Parenteau, M. Durand, S. Véronneau, A.-A. Lacombe, G. Morin, V. Guérin, B. Cecez, J. Gervais-Bird, C.-S. Koh, D. Brunelle, R. J. Wellinger, B. Chabot, S. Abou Elela, Deletion of many yeast introns reveals a minority of genes that require splicing for function. Mol. Biol. Cell 19, 1932–1941 (2008).

5. S. J. Gould, R. C. Lewontin, J. Maynard Smith, R. Holliday, The spandrels of San Marco and the Panglossian paradigm: a critique of the adaptationist programme. Proceedings of the Royal Society of London. Series B. Biological Sciences 205, 581–598 (1997).

6. E. Shoguchi, C. Shinzato, T. Kawashima, F. Gyoja, S. Mungpakdee, R. Koyanagi, T. Takeuchi, K. Hisata, M. Tanaka, M. Fujiwara, M. Hamada, A. Seidi, M. Fujie, T. Usami, H. Goto, S. Yamasaki, N. Arakaki, Y. Suzuki, S. Sugano, A. Toyoda, Y. Kuroki, A. Fujiyama, M. Medina, M. A. Coffroth, D. Bhattacharya, N. Satoh, Draft assembly of the Symbiodinium minutum nuclear genome reveals dinoflagellate gene structure. Curr. Biol. 23, 1399–1408 (2013).

7. S. W. Roy, W. Gilbert, The evolution of spliceosomal introns: patterns, puzzles and progress. Nat. Rev. Genet. 7, 211–221 (2006).

8. H. G. Morrison, A. G. McArthur, F. D. Gillin, S. B. Aley, R. D. Adam, G. J. Olsen, A. A. Best, W. Z. Cande, F. Chen, M. J. Cipriano, B. J. Davids, S. C. Dawson, H. G. Elmendorf, A. B. Hehl, M. E. Holder, S. M. Huse, U. U. Kim, E. Lasek-Nesselquist, G. Manning, A. Nigam, J. E. J. Nixon, D. Palm, N. E. Passamaneck, A. Prabhu, C. I. Reich, D. S. Reiner, J. Samuelson, S. G. Svard, M. L. Sogin, Genomic minimalism in the early diverging intestinal parasite Giardia lamblia. Science 317, 1921–1926 (2007).

9. P. Yenerall, L. Zhou, Identifying the mechanisms of intron gain: progress and trends. Biol. Direct 7, 29 (2012).

10. A. Z. Worden, J.-H. Lee, T. Mock, P. Rouzé, M. P. Simmons, A. L. Aerts, A. E. Allen, M. L. Cuvelier, E. Derelle, M. V. Everett, E. Foulon, J. Grimwood, H. Gundlach, B. Henrissat, C. Napoli, S. M. McDonald, M. S. Parker, S. Rombauts, A. Salamov, P. Von Dassow, J. H. Badger, P. M. Coutinho, E. Demir, I. Dubchak, C. Gentemann, W. Eikrem, J. E. Gready, U. John, W. Lanier, E. A. Lindquist, S. Lucas, K. F. X. Mayer, H. Moreau, F. Not, R. Otillar, O. Panaud, J. Pangilinan, I. Paulsen, B. Piegu, A. Poliakov, S. Robbens, J. Schmutz, E. Toulza, T. Wyss, A. Zelensky, K. Zhou, E. V. Armbrust, D. Bhattacharya, U. W. Goodenough, Y. Van de Peer, I. V. Grigoriev, Green evolution and dynamic adaptations revealed by genomes of the marine picoeukaryotes Micromonas. Science 324, 268–272 (2009).

11. J. T. Huff, D. Zilberman, S. W. Roy, Mechanism for DNA transposons to generate introns on genomic scales. Nature 538, 533–536 (2016).

12. L. Gozashti, S. W. Roy, B. Thornlow, A. Kramer, M. Ares Jr, R. Corbett-Detig, Transposable elements drive intron gain in diverse eukaryotes. Proc. Natl. Acad. Sci. U. S. A. 119, e2209766119 (2022).

13. S. W. Roy, L. Gozashti, B. A. Bowser, B. N. Weinstein, G. E. Larue, R. Corbett-Detig, Intron-rich dinoflagellate genomes driven by Introner transposable elements of unprecedented diversity. Curr. Biol. 33, 189–196.e4 (2023).

14. A. van der Burgt, E. Severing, P. J. G. M. de Wit, J. Collemare, Birth of new spliceosomal introns in fungi by multiplication of introner-like elements. Curr. Biol. 22, 1260–1265 (2012).

15. S. Farhat, P. Le, E. Kayal, B. Noel, E. Bigeard, E. Corre, F. Maumus, I. Florent, A. Alberti, J.-M. Aury, T. Barbeyron, R. Cai, C. Da Silva, B. Istace, K. Labadie, D. Marie, J. Mercier, T. Rukwavu, J. Szymczak, T. Tonon, C. Alves-de-Souza, P. Rouzé, Y. Van de Peer, P. Wincker, S. Rombauts, B. M. Porcel, L. Guillou, Rapid protein evolution, organellar reductions, and invasive intronic elements in the marine aerobic parasite dinoflagellate Amoebophrya spp. BMC Biol. 19, 1 (2021).

16. J. Collemare, A. van der Burgt, P. J. G. M. de Wit, At the origin of spliceosomal introns: Is multiplication of introner-like elements the main mechanism of intron gain in fungi? Commun. Integr. Biol. 6, e23147 (2013).

17. B. Verhelst, Y. Van de Peer, P. Rouzé, The complex intron landscape and massive intron invasion in a picoeukaryote provides insights into intron evolution. Genome Biol. Evol. 5, 2393–2401 (2013).

18. F. Denoeud, S. Henriet, S. Mungpakdee, J.-M. Aury, C. Da Silva, H. Brinkmann, J. Mikhaleva, L. C. Olsen, C. Jubin, C. Cañestro, J.-M. Bouquet, G. Danks, J. Poulain, C. Campsteijn, M. Adamski, I. Cross, F. Yadetie, M. Muffato, A. Louis, S. Butcher, G. Tsagkogeorga, A. Konrad, S. Singh, M. F. Jensen, E. Huynh Cong, H. Eikeseth-Otteraa, B. Noel, V. Anthouard, B. M. Porcel, R. Kachouri-Lafond, A. Nishino, M. Ugolini, P. Chourrout, H. Nishida, R. Aasland, S. Huzurbazar, E. Westhof, F. Delsuc, H. Lehrach, R. Reinhardt, J. Weissenbach, S. W. Roy, F. Artiguenave, J. H. Postlethwait, J. R. Manak, E. M. Thompson, O. Jaillon, L. Du Pasquier, P. Boudinot, D. A. Liberles, J.-N. Volff, H. Philippe, B. Lenhard, H. Roest Crollius, P. Wincker, D. Chourrout, Plasticity of animal genome architecture unmasked by rapid evolution of a pelagic tunicate. Science 330, 1381–1385 (2010).

19. T. Wicker, F. Sabot, A. Hua-Van, J. L. Bennetzen, P. Capy, B. Chalhoub, A. Flavell, P. Leroy, M. Morgante, O. Panaud, E. Paux, P. SanMiguel, A. H. Schulman, A unified classification system for eukaryotic transposable elements. Nat. Rev. Genet. 8, 973–982 (2007).

20. J. N. Wells, C. Feschotte, A field guide to eukaryotic transposable elements. Annu. Rev. Genet. 54, 539–561 (2020).

21. M. J. Curcio, K. M. Derbyshire, The outs and ins of transposition: from mu to kangaroo. Nat. Rev. Mol. Cell Biol. 4, 865–877 (2003).

22. M. Lynch, B. Walsh, The Origins of Genome Architecture (Sinauer Associates Sunderland, MA, 2007)vol. 98.

23. M. Krupovic, V. V. Dolja, E. V. Koonin, The virome of the last eukaryotic common ancestor and eukaryogenesis. Nat Microbiol 8, 1008–1017 (2023).

24. K. M. Devos, J. K. M. Brown, J. L. Bennetzen, Genome size reduction through illegitimate recombination counteracts genome expansion in Arabidopsis. Genome Res. 12, 1075–1079 (2002).

25. P. Capy, R. Vitalis, T. Langin, D. Higuet, C. Bazin, Relationships between transposable elements based upon the integrase-transposase domains: is there a common ancestor? J. Mol. Evol. 42, 359–368 (1996).

26. P. Capy, T. Langin, D. Higuet, P. Maurer, C. Bazin, “Do the integrases of LTR-retrotransposons and class II element transposases have a common ancestor?” in Evolution and Impact of Transposable Elements, P. Capy, Ed. (Springer Netherlands, Dordrecht, 1997), pp. 63–72.

27. H. Qiu, G. Cai, J. Luo, D. Bhattacharya, N. Zhang, Extensive horizontal gene transfers between plant pathogenic fungi. BMC Biol. 14, 41 (2016).

28. S. Schaack, C. Gilbert, C. Feschotte, Promiscuous DNA: horizontal transfer of transposable elements and why it matters for eukaryotic evolution. Trends Ecol. Evol. 25, 537–546 (2010).

29. S. Kumar, G. Stecher, M. Suleski, S. B. Hedges, TimeTree: A resource for timelines, timetrees, and divergence times. Mol. Biol. Evol. 34, 1812–1819 (2017).

30. M. Prieto, M. Wedin, Dating the diversification of the major lineages of Ascomycota (Fungi). PLoS One 8, e65576 (2013).

31. M. A. Van der Nest, E. T. Steenkamp, A. R. McTaggart, C. Trollip, T. Godlonton, E. Sauerman, D. Roodt, K. Naidoo, M. P. A. Coetzee, P. M. Wilken, M. J. Wingfield, B. D. Wingfield, Saprophytic and pathogenic fungi in the Ceratocystidaceae differ in their ability to metabolize plant-derived sucrose. BMC Evol. Biol. 15, 273 (2015).

32. S. Blair Hedges, S. Kumar, The Timetree of Life (OUP Oxford, 2009).

33. J. Copley, P. Tyler, M. Sheader Bramley, J. MURTONb, C. R. Germanb, Megafauna from sublittoral to abyssal depths along the Mid-Atlantic Ridge south of Iceland. Oceanol. Acta 19, 549–559 (1996).

34. J. Stephens, Atlantic sponges collected by the Scottish National Antarctic Expedition. Earth Environ. Sci. Trans. R. Soc. Edinb. 50 (1915).

35. M. Montresor, C. Lovejoy, L. Orsini, G. Procaccini, S. Roy, Bipolar distribution of the cyst-forming dinoflagellate Polarella glacialis. Polar Biol. 26, 186–194 (2003).

36. C. Rot, I. Goldfarb, M. Ilan, D. Huchon, Putative cross-kingdom horizontal gene transfer in sponge (Porifera) mitochondria. BMC Evol. Biol. 6, 71 (2006).

37. C. Conaco, P. Tsoulfas, O. Sakarya, A. Dolan, J. Werren, K. S. Kosik, Detection of prokaryotic genes in the Amphimedon queenslandica genome. PLoS One 11, e0151092 (2016).

38. J. H. Wisecaver, M. L. Brosnahan, J. D. Hackett, Horizontal gene transfer is a significant driver of gene innovation in dinoflagellates. Genome Biol. Evol. 5, 2368–2381 (2013).

39. X. Wang, X. Liu, Close ecological relationship among species facilitated horizontal transfer of retrotransposons. BMC Evol. Biol. 16, 201 (2016).

40. L. D. McDaniel, E. Young, J. Delaney, F. Ruhnau, K. B. Ritchie, J. H. Paul, High frequency of horizontal gene transfer in the oceans. Science 330, 50 (2010).

41. S. A. Widen, I. C. Bes, A. Koreshova, P. Pliota, D. Krogull, A. Burga, Virus-like transposons cross the species barrier and drive the evolution of genetic incompatibilities. Science 380, eade0705 (2023).

42. V. Loiseau, J. Peccoud, C. Bouzar, S. Guillier, J. Fan, G. Gueli Alletti, C. Meignin, E. A. Herniou, B. A. Federici, J. T. Wennmann, J. A. Jehle, R. Cordaux, C. Gilbert, Monitoring insect transposable elements in large double-stranded DNA viruses reveals host-to-virus and virus-to-virus transposition. Mol. Biol. Evol. 38, 3512–3530 (2021).

43. C. Gilbert, R. Cordaux, Viruses as vectors of horizontal transfer of genetic material in eukaryotes. Curr. Opin. Virol. 25, 16–22 (2017).

44. C. Gilbert, C. Feschotte, Horizontal acquisition of transposable elements and viral sequences: patterns and consequences. Curr. Opin. Genet. Dev. 49, 15–24 (2018).

45. N. A. T. Irwin, A. A. Pittis, T. A. Richards, P. J. Keeling, Systematic evaluation of horizontal gene transfer between eukaryotes and viruses. Nat Microbiol 7, 327–336 (2022).

46. T.-W. Sun, C.-L. Yang, T.-T. Kao, T.-H. Wang, M.-W. Lai, C. Ku, Host range and coding potential of eukaryotic giant viruses. Viruses 12 (2020).

47. F. Schulz, S. Roux, D. Paez-Espino, S. Jungbluth, D. A. Walsh, V. J. Denef, K. D. McMahon, K. T. Konstantinidis, E. A. Eloe-Fadrosh, N. C. Kyrpides, T. Woyke, Giant virus diversity and host interactions through global metagenomics. Nature 578, 432–436 (2020).

48. F. Schulz, N. Yutin, N. N. Ivanova, D. R. Ortega, T. K. Lee, J. Vierheilig, H. Daims, M. Horn, M. Wagner, G. J. Jensen, N. C. Kyrpides, E. V. Koonin, T. Woyke, Giant viruses with an expanded complement of translation system components. Science 356, 82–85 (2017).

49. L. Carmel, I. B. Rogozin, Y. I. Wolf, E. V. Koonin, Patterns of intron gain and conservation in eukaryotic genes. BMC Evol. Biol. 7, 192 (2007).

50. M. Csűrös, “Likely Scenarios of Intron Evolution” in Comparative Genomics (Springer Berlin Heidelberg, 2005), pp. 47–60.

51. H.-H. Zhang, J. Peccoud, M.-R.-X. Xu, X.-G. Zhang, C. Gilbert, Horizontal transfer and evolution of transposable elements in vertebrates. Nat. Commun. 11, 1362 (2020).

52. J. C. Slot, A. Rokas, Horizontal transfer of a large and highly toxic secondary metabolic gene cluster between fungi. Curr. Biol. 21, 134–139 (2011).

53. S. W. Roy, A. Fedorov, W. Gilbert, Large-scale comparison of intron positions in mammalian genes shows intron loss but no gain. Proc. Natl. Acad. Sci. U. S. A. 100, 7158–7162 (2003).

54. I. B. Rogozin, L. Carmel, M. Csuros, E. V. Koonin, Origin and evolution of spliceosomal introns. Biol. Direct 7, 11 (2012).

55. C. G. Sotero-Caio, R. N. Platt 2nd, A. Suh, D. A. Ray, Evolution and diversity of transposable elements in vertebrate genomes. Genome Biol. Evol. 9, 161–177 (2017).

56. J. O. Andersson, Lateral gene transfer in eukaryotes. Cell. Mol. Life Sci. 62, 1182–1197 (2005).

