## Supplementary Materials for "Horizontal transmission of functionally diverse transposons is a major source of new introns"

#### Materials and Methods

**Retrieving relevant genomic data**
We retrieved all annotated genomes available from the GenBank and RefSeq databases using NCBI's command line datasets tool (retrieved 08/28/2023). Metadata for all genomes accessed in this way are presented in table S1.

**Systematic candidate introner identification**
We used a previously developed pipeline for systematic introner identification (https://github.com/lgozasht/Introner-elements). A detailed description of this pipeline can be found in (*12*). Briefly, for each annotated genome, we extracted all introns, then performed an all vs. all blast (*57*) to search for highly similar introns. These introns were clustered based on sequence similarity (e-value < 1E−5), consistent with signatures expected for recently active transposable elements. This clustering produced families of candidate introners.

Since several alternative reasons could account for sequence similarity between introns, we then performed several filtering steps. First, we filtered candidate introners which displayed sequence similarity due to secondary insertion of transposable elements into preexisting introns. To do this, we required that sequence similarity extended to the exon-intron boundaries, but not into the exons for each intron in the expected orientation (5' regions aligned with 5' regions and 3' regions aligned with 3' regions across candidate introners in a given family). Second, we removed introns which displayed sequence similarity as a result of whole gene duplication. To do this, we filtered candidate introners which displayed sequence similarity extending into exons, assuming that duplication of an entire intron without any flanking exonic sequence is unlikely. While this filter removed the vast majority of false positives due to paralogous gene duplication, our previous work suggests that rapid exonic evolution (or simple chance substitution) can lead to paralogous sequences being retained in some cases (such as for the large, fast-evolving var gene families of *Plasmodium* species) (*12*). To filter the remaining false positives caused by introns in paralogous genes, we translated all introner-containing genes and used diamond (version 0.9.24) (*58*) (e-value < 1E-20) to identify cases of sequence similarity between encoded proteins (paralogy groups) in each species. We retained all introner families with at least 4 sequences from genes in different paralogy groups that passed these filters.

**Manual curation of candidate introners**
Manual curation is essential for validating candidates introner and transposable element models more generally (*12*, *59*). We performed several steps of manual inspection to validate candidate introners. First, we generated multiple sequence alignments for each candidate introner family using MAFFT (*60*) and viewed alignments using Aliview (*61*). We manually checked multiple sequence alignments to confirm that introner homology extends to near to the intron edges but not far into flanking exons, and that homology boundaries are consistent across all introners in a given family. In doing so, we manually removed spurious introners resulting from gene duplications, which may have inadvertently passed our systematic filters (above). We also inspected introner sequences for low complexity regions and removed introners for which

sequence homology was primarily driven by simple repeats or satellites. Finally, we manually inspected introner sequences for signatures of spurious intron annotations. We checked that 5' and 3' splice sites were largely consistent for all introners in a given subfamily and further interrogated introner families which primarily showed inconsistent or unfamiliar splice sites. Canonical and noncanonical 5' splices sites for major and minor introns generally include GT, GC, GA and AT, whereas 3' splice sites include AG and AC (*62*, *63*). Due to the possibility of annotation errors, for introner families with primarily non-canonical splice sites, we manually inspected splice junctions in NCBI's genome data viewer to ensure correct splicing of introners and expression of introner-containing genes based on transcript or RNA-seq alignments. We discarded introner families which did not meet these requirements. We constructed consensus sequences and calculated Kimura divergence from the consensus for introner copies using RepeatModeler's utility tool, Refiner (*64*).

**Introner TE functional classification**

We classified introners as they relate to transposable elements using a combination of systematic and manual approaches. First, we ran RepeatClassifier (*64*) on introner consensus sequences using all curated TE models available from the DFAM (*65*) and RepBase (*66*) databases. RepeatClassifier is a homology based approach that prioritizes accuracy rather than sensitivity. Since many species in our data are divergent from those with curated TE models, we also classified introners *de novo* by annotating structural features and functional domains that might be consistent with transposition. Indeed, various transposable elements display structural features such as long terminal repeats or terminal inverted repeats, and the presence of certain protein domains, such as a reverse transcriptase, can be used to classify transposable elements.

We used TE-AID (*67*) to annotate and visualize introner sequence structures. TE-AID produces four different types of plots which aid in TE classification (see fig. S8 for example). First, TE-AID retrieves all prospective copies for a given TE family by blasting the consensus sequence at the respective reference genome and plots divergence from the consensus for all fragmented and full length copies. Then, it plots coverage with respect to the consensus. This especially aids with classifying LTR retrotransposons since LTRs at element edges often exhibit much higher copy numbers than interior regions due to frequent ectopic recombination between LTRs resulting in the interior region's removal (*24*). TE-AID also produced selv-v-self dotplots for TE-consensus sequences, allowing us to assess low complexity regions as well as hallmark features of different TE mechanisms such as LTRs and TIRs. Finally, TE-AID produces a plot showing the locations of open reading frames (ORFs) within each TE-consensus as well as their homology to known TEs.

We also sought to identify functional machinery and protein features within introner sequences. To do this, we first predicted and translated all possible ORFs in introner sequences using orfm (*68*). Then, we used interproscan (version 5.65-97.0) (*69*, *70*) to identify functional protein-coding domains and features in all introner sequences. We also separately ran hmmer (*71*) on ORFs from introner consensus sequences to scan for TE-related functional domains using all available models from the PFAM and Gypsy databases (*72*, *73*). Since specific protein machinery is associated with different known TE mechanisms, these results provided evidence for introner TE classifications.

For introners with no clear classification based on these methods, we ran MCHelper (*74*) to further extend and refine introner consensus sequences. MCHelper also automatically identifies structural features and functional domains in refined consensus sequences. Due to the challenges associated with automated transposable element extension and refinement (*67*), we hypothesized that some introner consensus sequences produced by Refiner may have been overextended or may exhibit other problems that inhibit automated feature discovery and classification. Thus, we ran MCHelper using three different inputs for each introner: (1) using the entire consensus, (2) using the first half of the consensus as input (and again using the second half if the first half yielded no results) and (3) using the middle 100 bp as input (to account for consensus overextension). This allowed us to classify additional introners with previously ambiguous classifications (table S2). Since many introners are nonautonomous and are challenging to classify due to their lack of functional transposition machinery, we also retrieved HMMs for conserved termini of known transposable elements available from DFAM. We used nhmmer (*75*) to search for these conserved termini in introner families, filtering for hits with E<0.001. We classified additional introners based on the presence of these conserved termini at expected positions in introner consensus sequences.

**Validating novel introner-containing taxa**

We performed additional steps to validate the presence and correct assembly of introners in several divergent taxa previously not known to contain them, which include *Strongylocentrotus purpuratus* (purple sea urchin), *Panicum virgatum* (switchgrass), and *Styela clava* (tunicate). While this list is not exhaustive, it captures the broad diversity of newly-discovered introner containing species. Generally, we checked for splicing of the introners based on mapped RNA-Seq reads, and looked at orthologous genes in a closely related species and a more distant relative to determine if the insertion caused intron gain. For *S. purpuratus* and *S. clava*, we utilized the genome browser available on NCBI for RefSeq genomes to view mapped RNA-Seq reads and orthologous genes, and used Phytozome (*76*) to do this for *P. virgatum*. For *S. purpuratus*, we additionally utilized available PacBio datasets to check that mapped long reads spanned the introner regions and did not indicate a deletion of the putative introner (fig. S1-S7). *S. purpuratus* long read accessions used for mapping and the number of reads that mapped from each are listed in table S11. In all cases, we were able to successfully validate the presence of intron-generating introners through long read coverage that extended well into the flanking portions of the genome from introner insertions.

**Searching for horizontal gene transfer**

To search for possible HGT of introners, we performed blastn searches (*57*) for each introner consensus sequence against the NCBI nucleotide and RefSeq reference genome databases (accessed 05/15/2024) (*77*). Then, we filtered for hits with e-values <0.0001, percent sequence ID > 70 and coverage > 60% in genera other than the source species'. These results are displayed in table S4. We emphasize that these cutoffs are conservative and we expect to only find the most recently horizontally transferred elements between highly genetically divergent host species about which we can be the most confident. Furthermore, many genera contain exceptionally diverse species, and this simple approach may exclude HGT that occurs between distantly-related species that nonetheless share a single genus. Then, we manually inspected results to filter out false positives resulting from conserved regions of transposons (e.g. regions of the RT domain in retrotransposons (*78*)) as well as low-quality subject sequences.

These efforts identified 8 candidate instances of HGT of recently active introners (table S5). Together, introner HGT events involve 11/201 introner-containing lineages (5.5% of surveyed species). Some HGT events (4/9) involve introners actively generating introns in both lineages, whereas for other HGT events (5/9), introners are only actively generating introns in one lineage and not the other. For example, the ascomycete fungi, *Xylaria* and *Lasiodiplodia* (diverged 350 MYA) and *Alternaria* and *Parastagonospora* (diverged 134 MYA) both display highly similar active introner families driving ongoing intron gain (*29–31*). In contrast, introners in the lichen *Amylostereum chailletii* display high sequence similarity to sequences in *Stereum hirsutum* (112 MY diverged; table S4-S5). However, we do not observe introner-mediated intron gain in *Stereum hirsutum* (table S2). When evaluating evidence of HGT, we retrieved homologous sequences from distantly related species iteratively using NCBI's entrez tool through BioPython (*79*).

We performed additional confirmation of HGT between the dinoflagellate, *Polarella glacialis*, and the glass sponge, *Aphrocallistes beatrix*, as this represents the HGT event between the most distantly related eukaryotic lineages that we detected using this approach. The correct assembly of introners in *P. glacialis* was determined by mapping long reads, as described above for confirming bacterial insertions. We also utilized this method to check for the correct assembly of regions in *A. beatrix* with homology to *P. glacialis* introner sequences, as found by the blastn searches. This allowed us to successfully validate the presence of introners in *P. glacialis*, as well as their non-intron generating counterparts in *A. beatrix* (fig. S12, tables S8-S9). We used this same approach to validate HGT of introners and non-introner-generating homologs between the diatom, *Thalassiosira oceanica* and the leech, *Piscicola geometra* (fig. S15; table S12).

**Phylogenetic Inference**

For instances of HGT, we attempted phylogenetic reconstruction to estimate the approximate time of transfer and to investigate patterns of transmission within and across the host genome. To do this we performed multiple sequence alignment of introner families (or regions homologous to introners in the source species) using MAFFT with the --adjustdirectionaccurately flag. Then, we calculated mean percent identity for all interspecies introner comparisons.

Maximum likelihood phylogenies were inferred from the ascomycete introner alignments with W-IQ-Tree (*80*), using the ModelFinder option, 0.5 perturbation strength, 100 unsuccessful iterations till stop, generalized midpoint root optimization, and 100 standard bootstrap replicates.

To isolate the informative sites in the highly polymorphic dinoflagellate and sponge introner sequence alignments, we used ClipKIT (*81*) to trim regions with gaps in greater than 90% of sequences and retain only parsimony-informative and constant sites (mode: kpic-gappy). Next, we inferred maximum likelihood phylogenies from the dinoflagellate and sponge introner elements with IQ-Tree (*82*) using the ModelFinder option and 100 standard bootstrap replicates.

We estimated time-scaled phylogenies with TreeTime (*83*), specifying sampling dates from NCBI and substitution rates from the literature. For ascomycete fungi, neutral rates range from 1e−8 to 1e−9 substitutions per site per year (*84*, *85*). Consistent with this value, the substitution rate for full-length LTR retrotransposons in fungi has been estimated to be 1.3e-8 substitutions per site per year (*86*). We used a substitution rate of 1.3e-8 for our ascomycete introner divergence time estimates. While substitution rates have not yet been measured for any sponges, they have been measured for a wide range of invertebrate taxa. The estimated genome-wide neutral substitution rate for *Alpheus* snapping shrimp is 2.64e−9 substitutions per site year (*87*) and the average rate across insects and molluscs is estimated to be 4.40e-9 per site per year (*88*). In diatoms, the base substitution mutation rate per site per generation has been measured to be 4.77e−10 and approximately 1.32 generations occur per day (*89*). This short generation time equates to 2.30e-7 substitutions per site per year (4.77e−10 subs/site/generation * 365 days/year * 1.32 generations/day). From these rates, we used the lowest (2.64e-9) for our introner divergence date estimates because it is the most conservative.

We caution that due to the exceptionally short lengths of sequences considered that there is substantial phylogenetic uncertainty in the resulting trees (fig. S10-S11, S13 contain bootstrapped phylogenies). Nonetheless, the overarching patterns are readily apparent and support our interpretation.

To compare introner divergence time estimates to organismal divergence times, we obtained dated species trees from the TimeTree database (*29*) for each of the focal taxonomic groups containing introner HGT events. The *Polarella* clade was not represented in TimeTree. So we identified *Symbiodinium* as another dinoflagellate in the same order (Suessiales), which was available. Given the deep split between *Symbiodinium* and its sister group, the Gymnodiniales order, the topology of the tree is likely the same for *Symbiodinium* as it would be for *Polarella*, if it had been sampled. Multiple sequence alignments underlying introner phylogenies in Figure 3B,C,E are available at https://github.com/lgozasht/Introner-elements/tree/main/msa.

**Searching for introners in genes of viral origin**

To identify introners in genes of viral origin, we retrieved all introner-containing proteins from NCBI and performed tblastn (*57*) searches against the Reference Viral DataBase (RVDB v28.0; accessed May 2, 2024) (*90*). We then filtered for hits with evalues < 0.0001. This revealed several high-confidence instances of introner-containing genes of viral origin (table S10). To test for overrepresentation of introner-containing genes among genes with homology to viral proteins, we randomly resampled genes from each introner-containing genome (N=number of introner-containing genes). We performed tblastn searches against RVDB with these randomly sampled genes to derive an expected distribution of genes showing homology to viral proteins in each species. Then, we performed two statistical tests for enrichment. First, we compared the observed distribution of candidate introner-containing viral-derived genes to an expected distribution aggregated across species using a nonparametric MWU test ($P<0.00001$). Second, we performed a two-way binomial test comparing the number of species with a greater number of introner-containin genes represented among viral proteins than expected ($P<0.0001$). To search for possible overrepresentation of introner-containing genes among specific viral lineages, we first retrieved NCBI lineage taxonomic data for each viral accession in RVDB using ete3 (*91*, *92*). This revealed that ~10% of introner-containing genes with homology to viral proteins mapped to proteins in klosneuviral genomes. To test for overrepresentation of introner-containing proteins among klosneuviral sequences, we performed a two-way binomial test where the null expectation was the proportion of nucleotides in RVDB represented by klosneuviral sequences.

**Figure 1 cladogram**
To generate a cladogram of eukaryotic lineages, we first retrieved NCBI taxonomy IDs for the phylum and order corresponding to each considered species. Then, we assembled a list of nonredundant phyla from these results. For eukaryotic lineages with no taxonomic ID for phylum, we used order instead. Then, we submitted this list of nonredundant taxonomy IDs to Interactive Tree of Life (ITOL), which produced a cladogram in newick format (*93*). The resulting tree was manipulated using ete3 (*92*) and visualized using toytree (*94*).

#### Supplemental Text

**Bacterial HGT in *Polarella glacialis***
We found evidence of HGT between bacteria and *Polarella glacialis* strain CCMP2088 in introner regions, providing additional evidence of inter-domain transfer in dinoflagellates. To confirm the bacterial insertions in *Polarella glacialis* strain CCMP2088, we first used a blastn search against the NCBI bacterial nt database to identify hits to bacterial sequences in the introner regions. We then retrieved *P. glacialis* PacBio long reads from the NCBI SRA (list of accessions and number of mapped reads in table S13), mapped them back to the reference genome using Minimap2 (*95*), and calculated the coverage at each introner position using bedtools multicov (*96*). Finally, we filtered, sorted, and merged bam files using samtools, and viewed the mapped reads, introner positions, and bacterial blast hit positions using the Integrated Genomics Viewer (IGV) (*97*). Introner sequences were confirmed to be true insertions when long reads spanned the entire introner and mapped past the introner boundaries on both sides. Examples are shown in fig. S14.

### Supplemental Figures

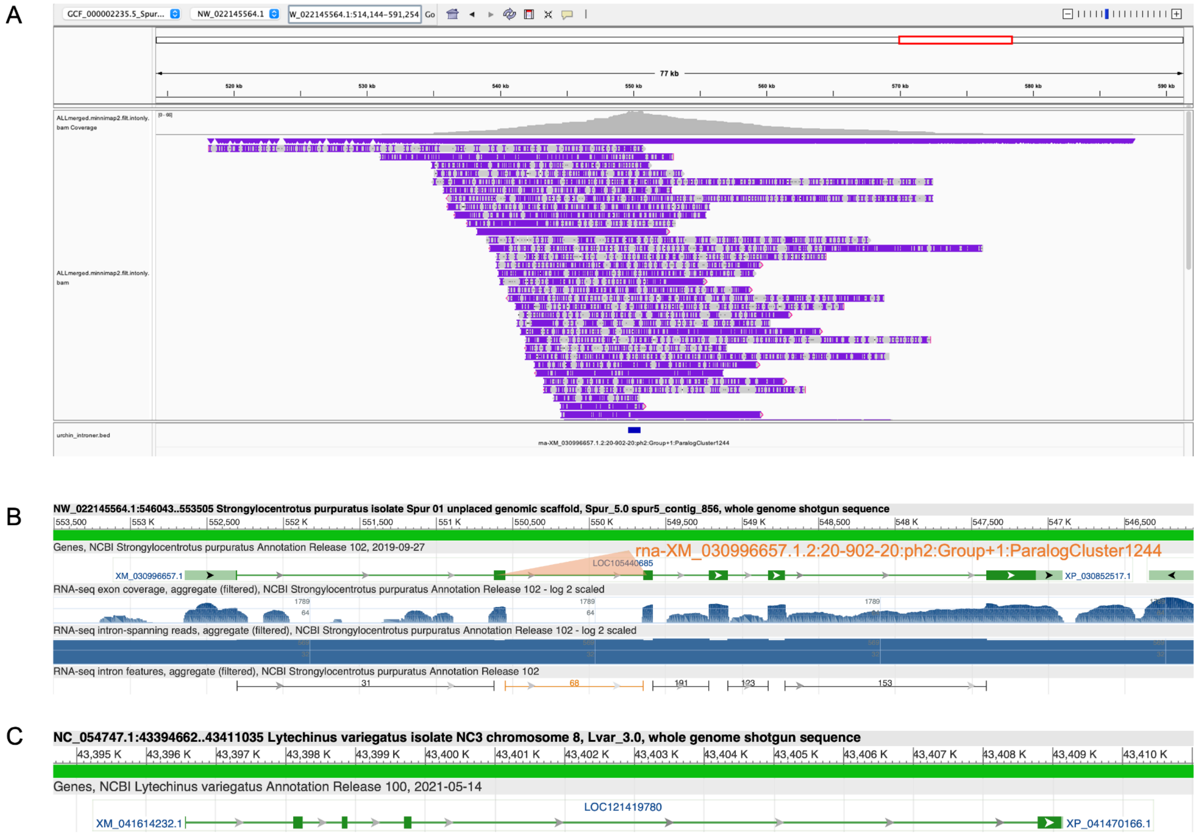


**fig. S1:** Verification of an introner (rna-XM_030996657.1.2:20-902-20:ph2:Group+1:ParalogCluster1244 located at NW_022145564.1:550569-549628) in *Strongylocentrotus purpuratus* (purple sea urchin, GCF_000002235.5). **(A)** Mapped long reads (PacBio) span the introner region (only reads overlapping the introner are shown). **(B)** Mapped RNA-seq coverage shows clear breaks, indicating splicing of the intron. **(C)** An ortholog in a close relative species (*Lytechinus variegatus*, green sea urchin) contains one less intron, supporting intron gain via the introner in *S. purpuratus*.

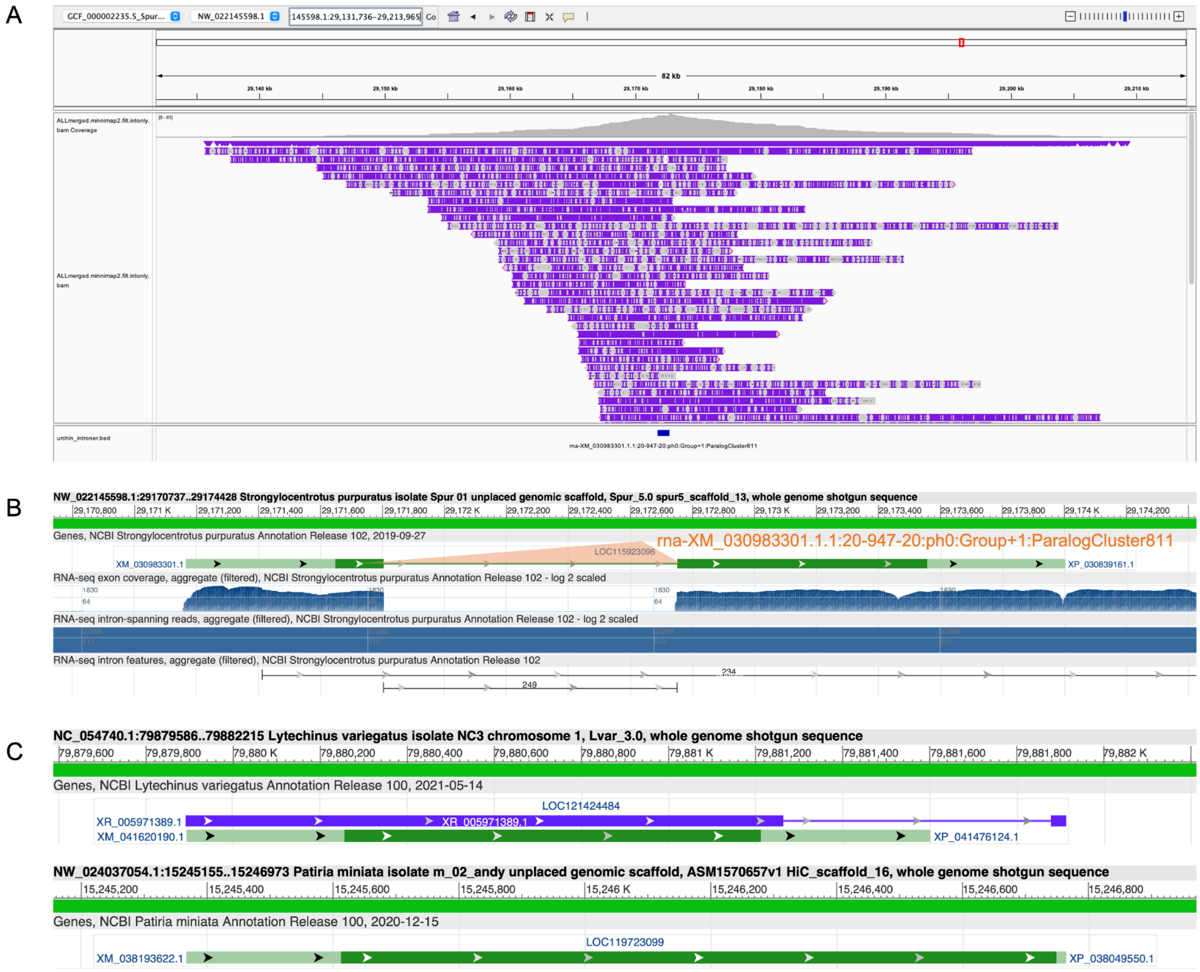


**fig. S2:** Verification of an introner (rna-XM_030983301.1.1:20-947-20:ph0:Group+1:ParalogCluster811 located at NW_022145598.1:29171784-29172770) in *Strongylocentrotus purpuratus* (purple sea urchin, GCF_000002235.5). **(A)** Mapped long reads (PacBio) span the introner region (only reads overlapping the introner are shown). **(B)** Mapped RNA-seq coverage shows clear breaks, indicating splicing of the intron. **(C)** Orthologs in a close relative species (*Lytechinus variegatus*, green sea urchin) and distantly related species (*Patiria miniata*) do not contain introns, supporting intron gain via the introner in *S. purpuratus*.

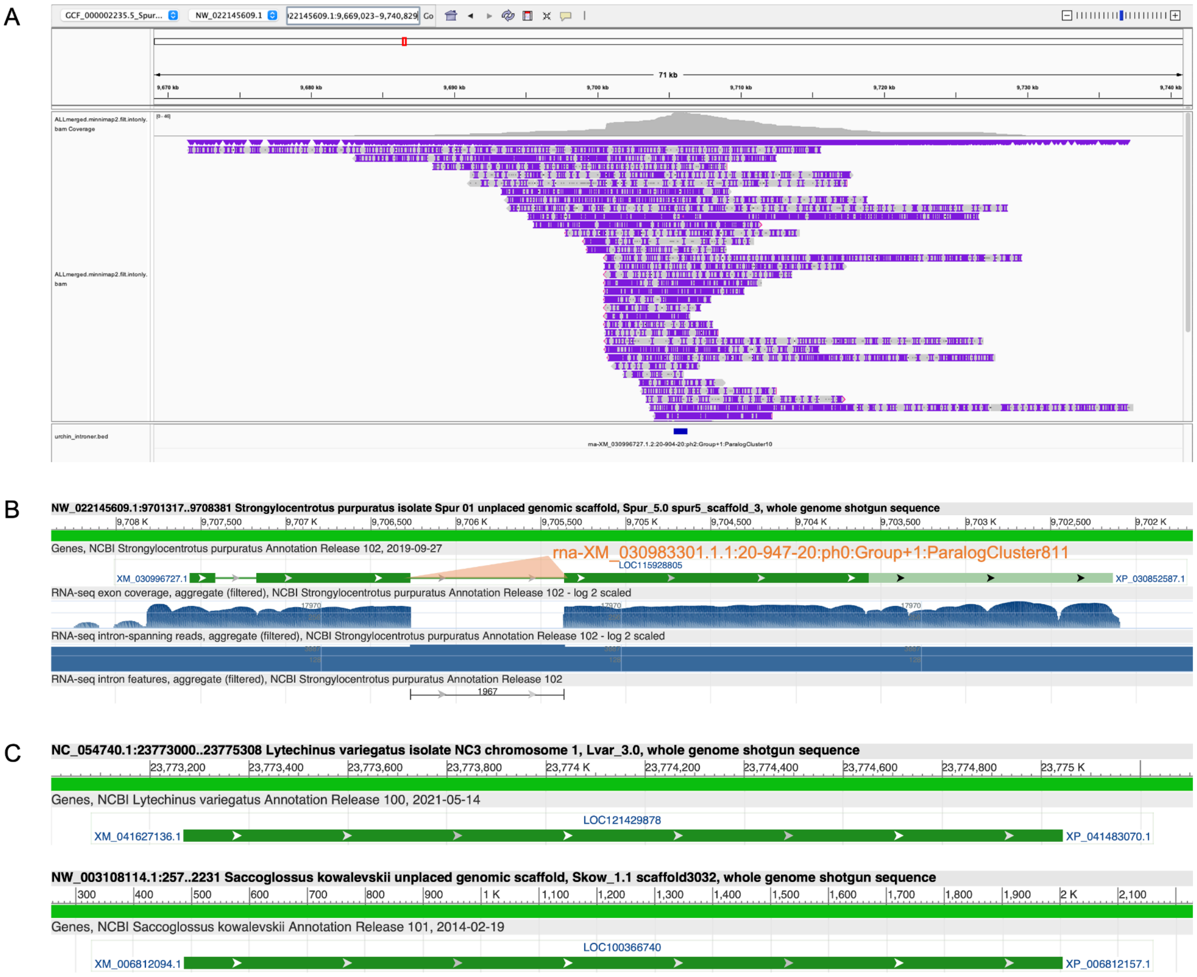


**fig. S3:** Verification of an introner (rna-XM_030996727.1.2:20-904-20:ph2:Group+1:ParalogCluster10 located at NW_022145609.1:9706287-9705344) in *Strongylocentrotus purpuratus* (purple sea urchin, GCF_000002235.5). **(A)** Mapped long reads (PacBio) span the introner region (only reads overlapping the introner are shown). **(B)** Mapped RNA-seq coverage shows clear breaks, indicating splicing of the intron. **(C)** Orthologs in a close relative species (*Lytechinus variegatus*, green sea urchin) and distantly related species (*Saccoglossus kowalevskii*) do not contain introns, supporting intron gain via the introner in *S. purpuratus*.

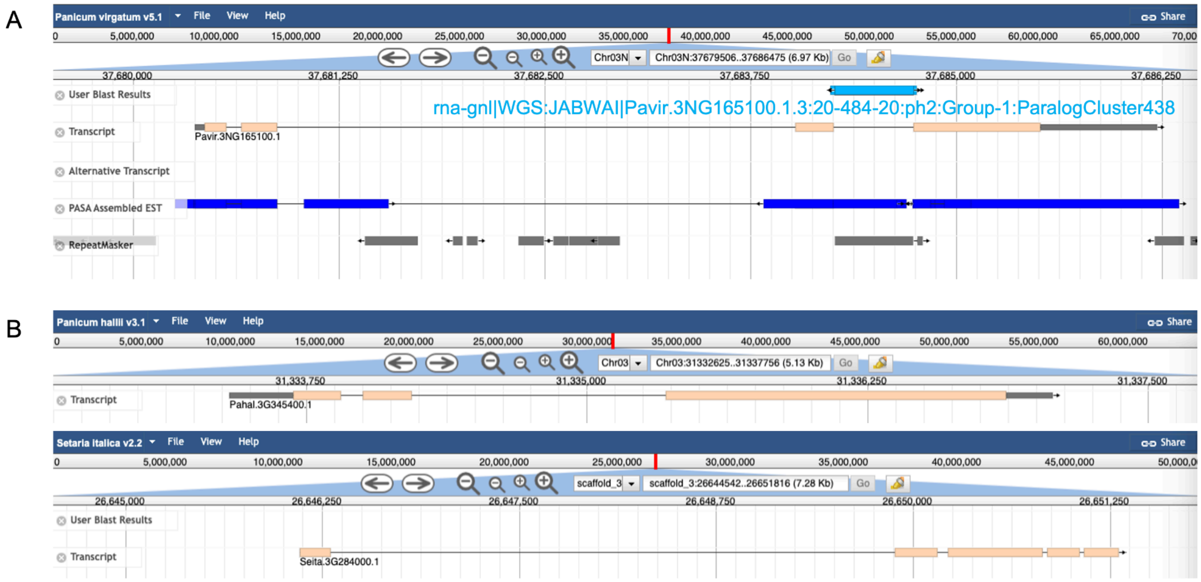


**fig. S4:** Verification of an introner (rna-gnl|WGS:JABWAI|Pavir.3NG165100.1.3:20-484-20:ph2:Group-1:ParalogCluster438 located at CM029042.1:37686220-37686704) in *Panicum virgatum* (switchgrass, GCA_016808335.1). **(A)** Transcripts based on RNA-seq reads indicate splicing of the intron. **(B)** An ortholog in a close relative species (*Panicum hallii*) and distantly related species (*Setaria italica*) contain no intron at the introner position, supporting intron gain via the introner in *P. virgatum*.

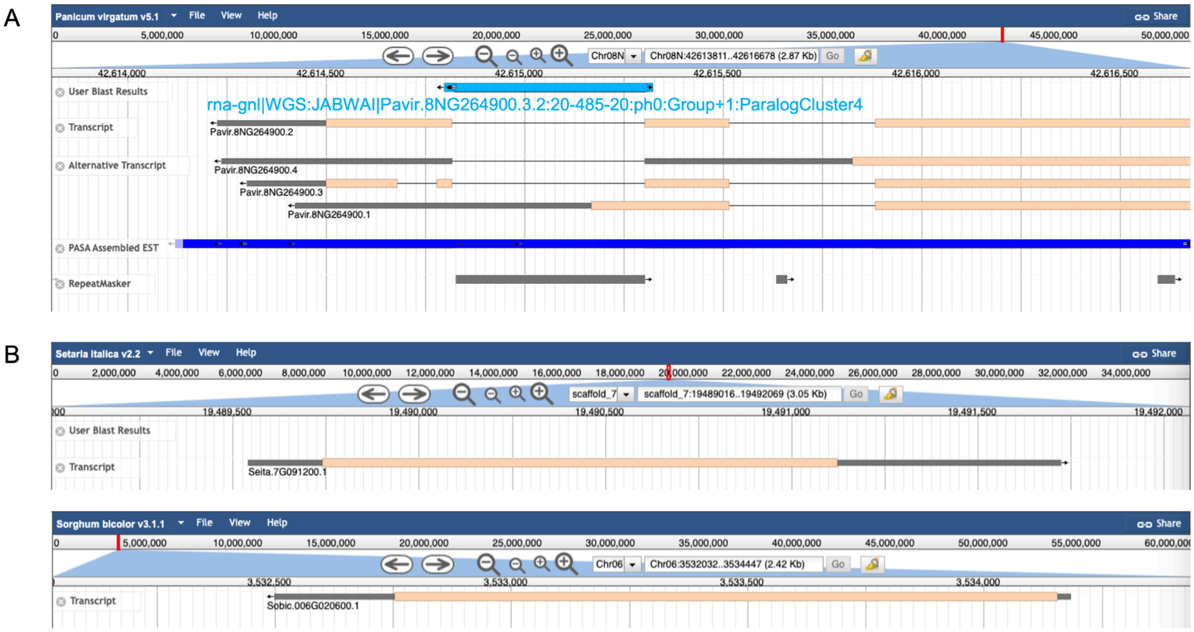


**fig. S5:** Verification of an introner (rna-gnl|WGS:JABWAI|Pavir.8NG264900.3.2:20-485-20:ph0:Group+1:ParalogCluster422 located at CM029052.1:42616768-42617253) in *Panicum virgatum* (switchgrass, GCA_016808335.1). **(A)** Transcripts based on RNA-seq reads indicate splicing of the intron. **(B)** Orthologs in relative species (*Setaria italica* and *Sorghum bicolor*) contain no introns, supporting intron gain via the introner in *P. virgatum*.

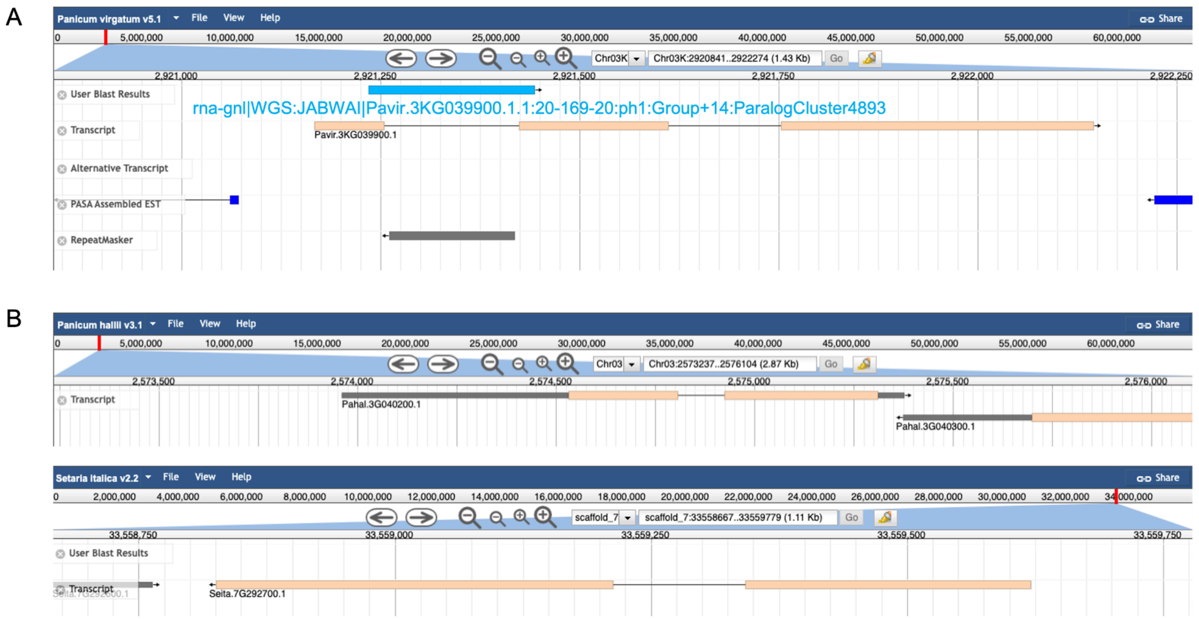


**fig. S6:** Verification of an introner (rna-gnl|WGS:JABWAI|Pavir.3KG039900.1.1:20-169-20:ph1:Group+14:ParalogCluster4893 located at CM029041.1:2922145-2922314) in *Panicum virgatum* (switchgrass, GCA_016808335.1). **(A)** Transcripts based on RNA-seq reads indicate splicing of the intron. **(B)** An ortholog in a close relative species (*Panicum hallii*) and distantly related species (*Setaria italica*) contain one less intron, supporting intron gain via the introner in *P. virgatum*.

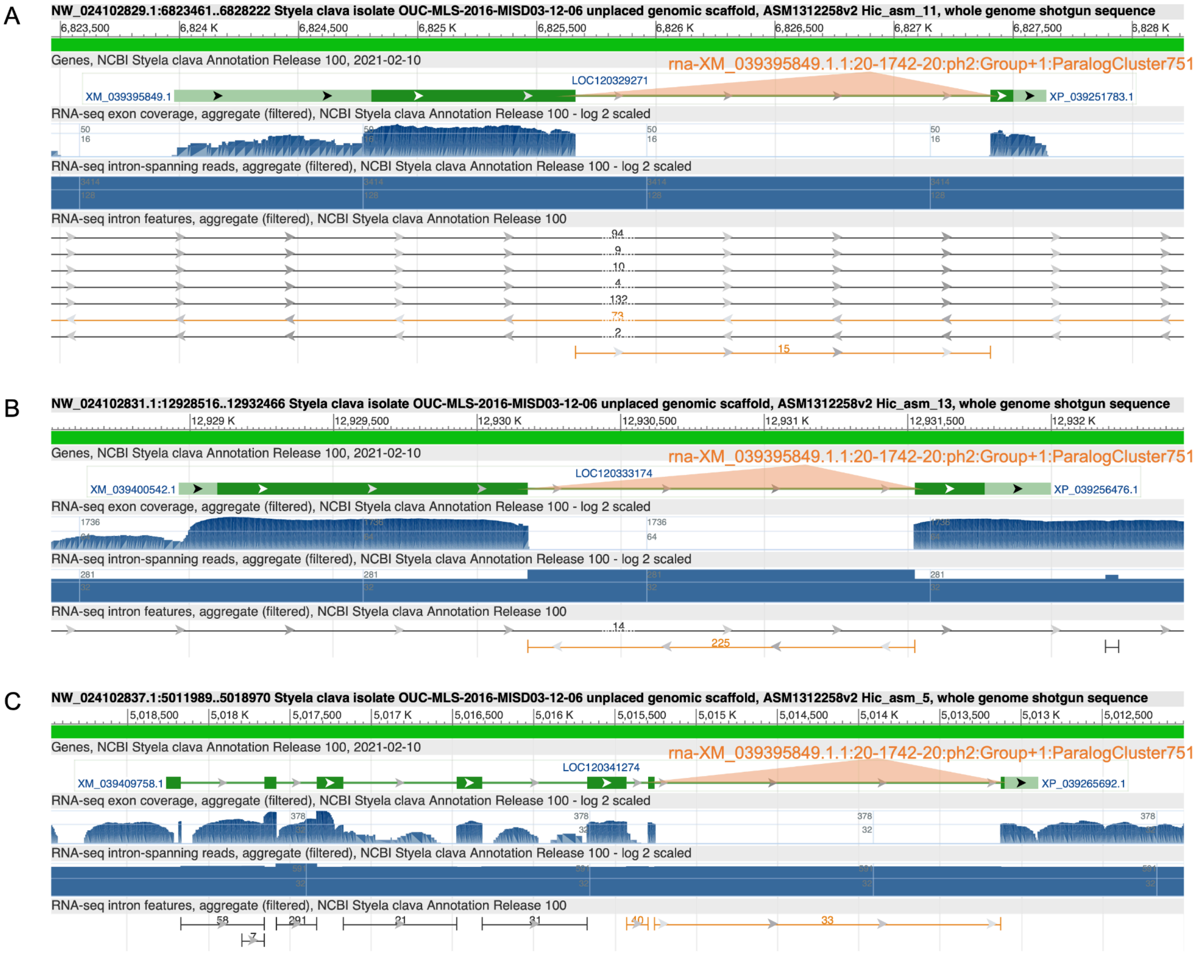


**fig. S7:** Verification of introners in *Styela clava* (tunicate, GCF_013122585.1). Mapped RNA-seq coverage shows clear breaks, indicating splicing of the intron in **(A)** rna-XM_039395849.1.1:20-1742-20:ph2:Group+1:ParalogCluster751 located at NW_024102829.1:6825643-6827424, **(B)** rna-XM_039400542.1.1:20-1348-20:ph2:Group-1:ParalogCluster544 located at NW_024102831.1:12930157-12931544, and **(C)** rna-XM_039409758.1.6:20-2131-20:ph1:Group+1:ParalogCluster221 located at NW_024102837.1:5015276-5013106.

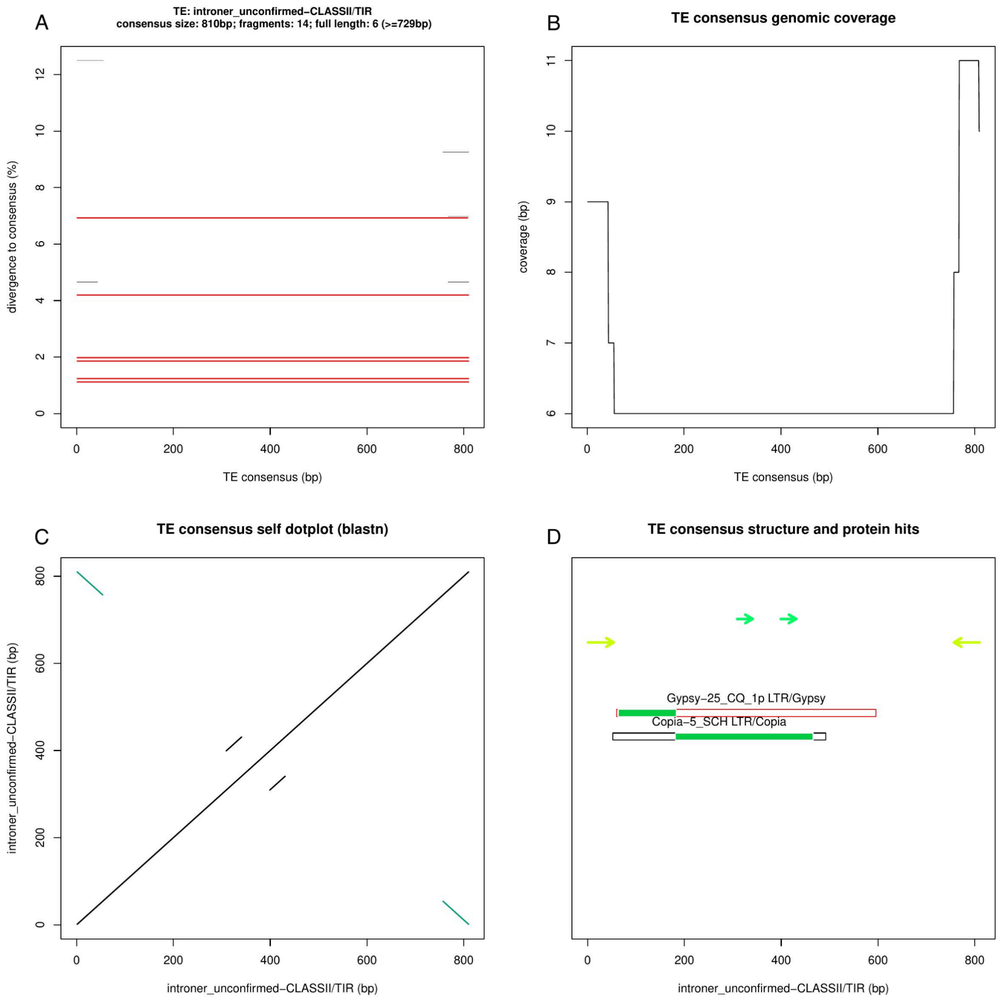


**fig. S8:** Structure and homology to known TEs for *Amoebophrya sp.* A120 family 38 introners. (**A**) Divergence from the consensus for introner copies. (**B**) Genomic coverage with respect to introner consensus. (**C)** Dotplot displaying terminal inverted repeats (TIRs) at introner edges. (**D**) Structural features and homology to known TEs for introner ORFs. Black boxes represent introner ORFs and green shading denotes regions of homology to known TEs. TIRs are shown as inverted arrows, and direct repeats are shown with collinear arrows. Plots produced with TE-AID (*67*).

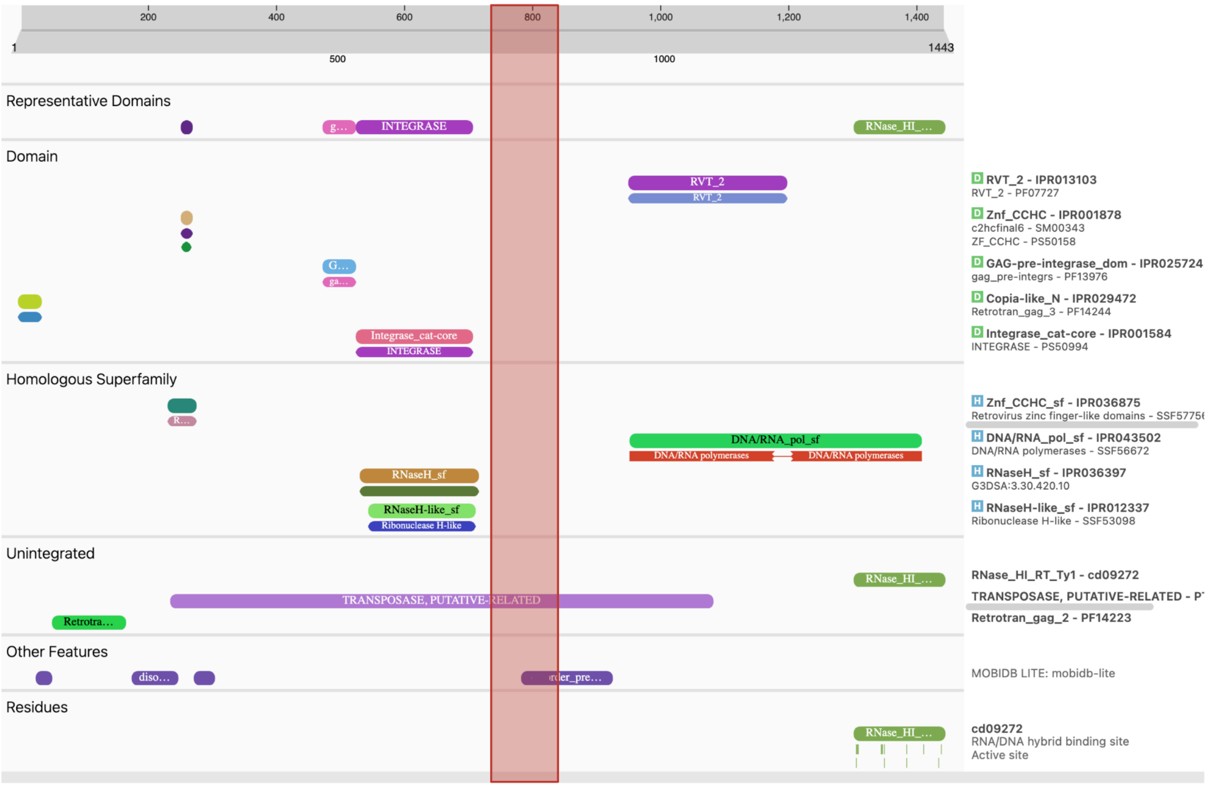


**fig. S9:** Interproscan results for Copia-5_SCH-I (RepBase), which shows homology to *Amoebophrya sp.* A120 family 38. The homologous region to *Amoebophrya sp.* A120 family 38 is highlighted in red. Figure produced using the Interpro web browser (*70*).

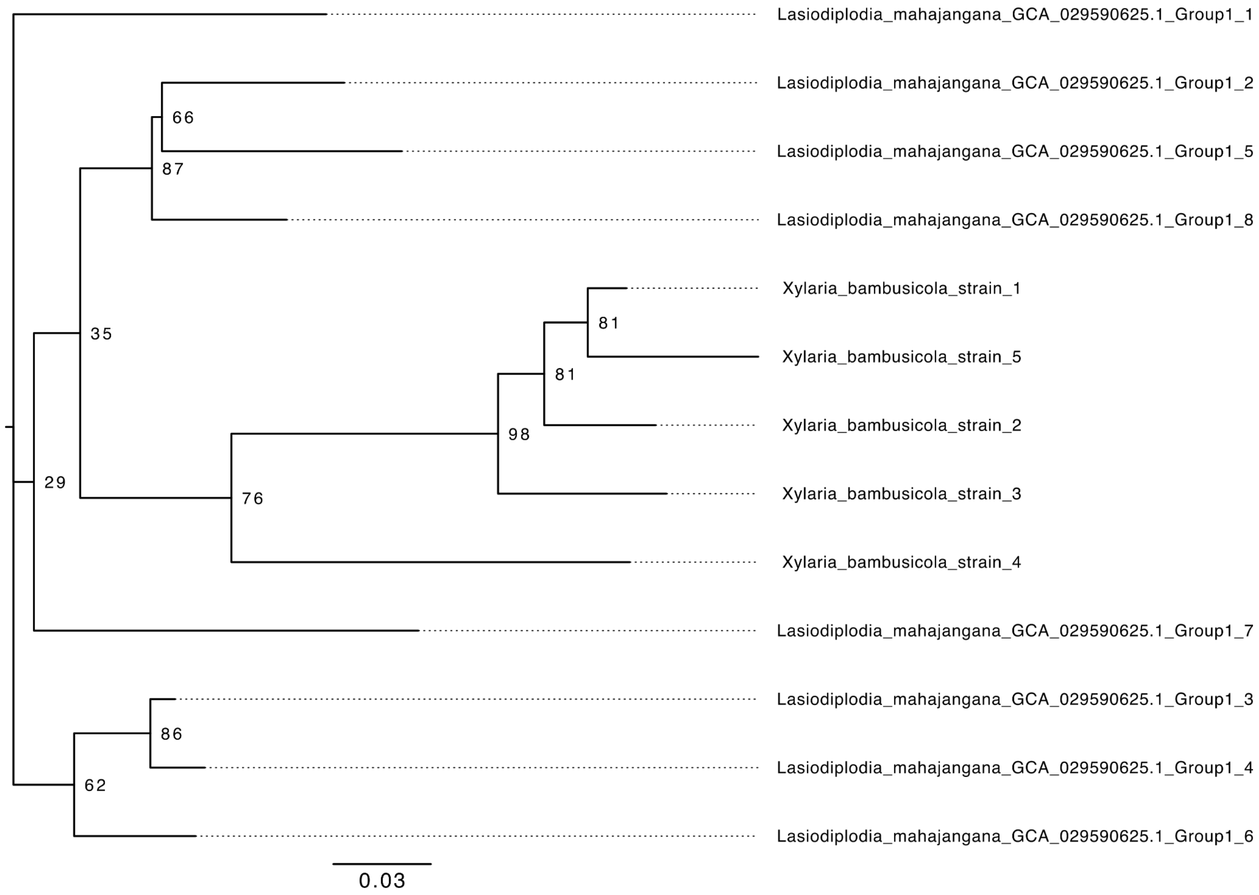


**fig. S10:** Maximum likelihood phylogeny for the introner found in *Lasiodiplodia mahajangana* and *Xylaria bambusicola* fungi, rooted at the midpoint. Bootstrap values out of 100 are indicated at the nodes. Scale bar measures substitutions per site.

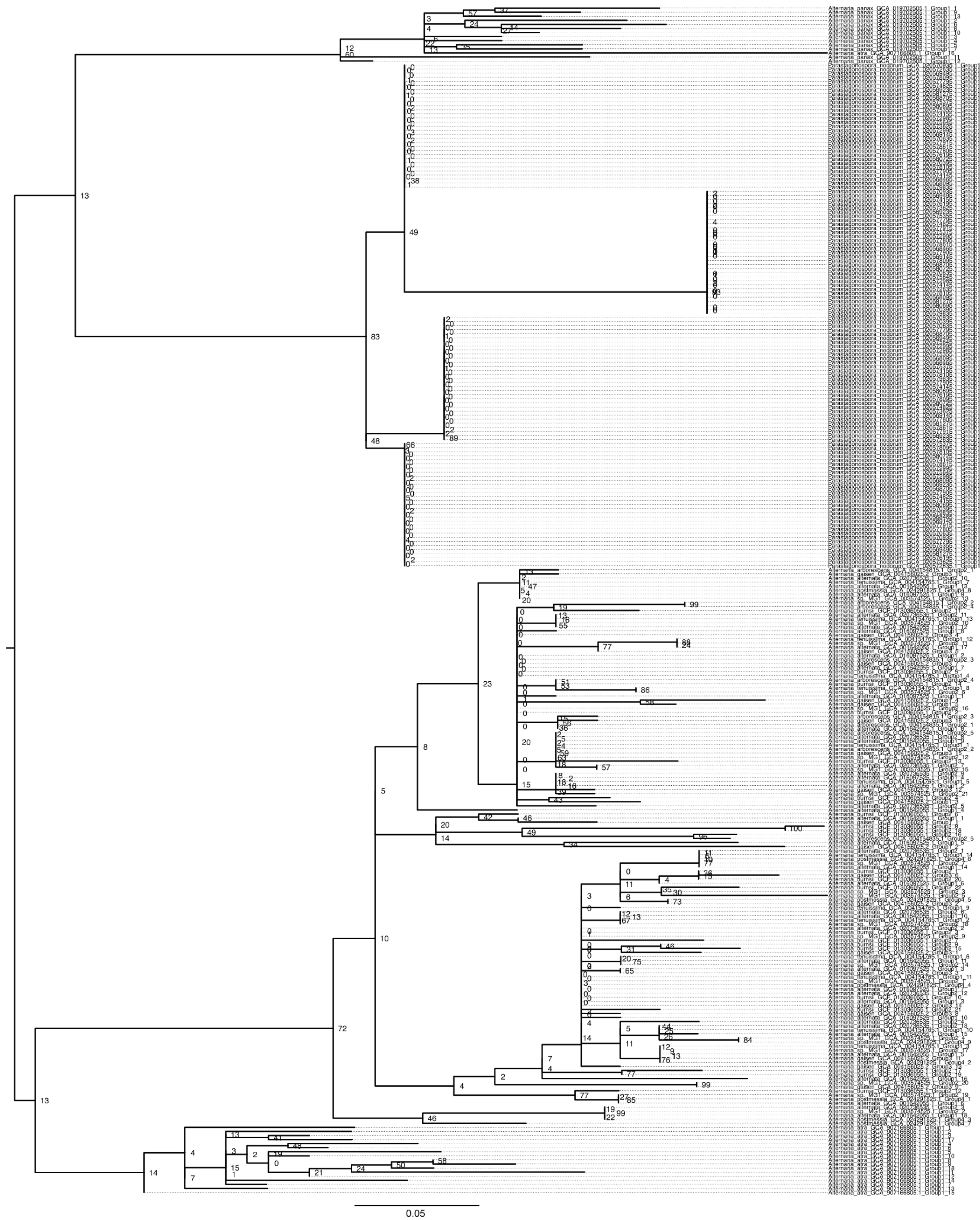


**fig. S11:** Maximum likelihood phylogeny for the introner found in *Parastagonospora nodorum* and *Alternaria spp*. fungi, rooted at the midpoint. Bootstrap values out of 100 are indicated at the nodes. Scale bar measures substitutions per site.

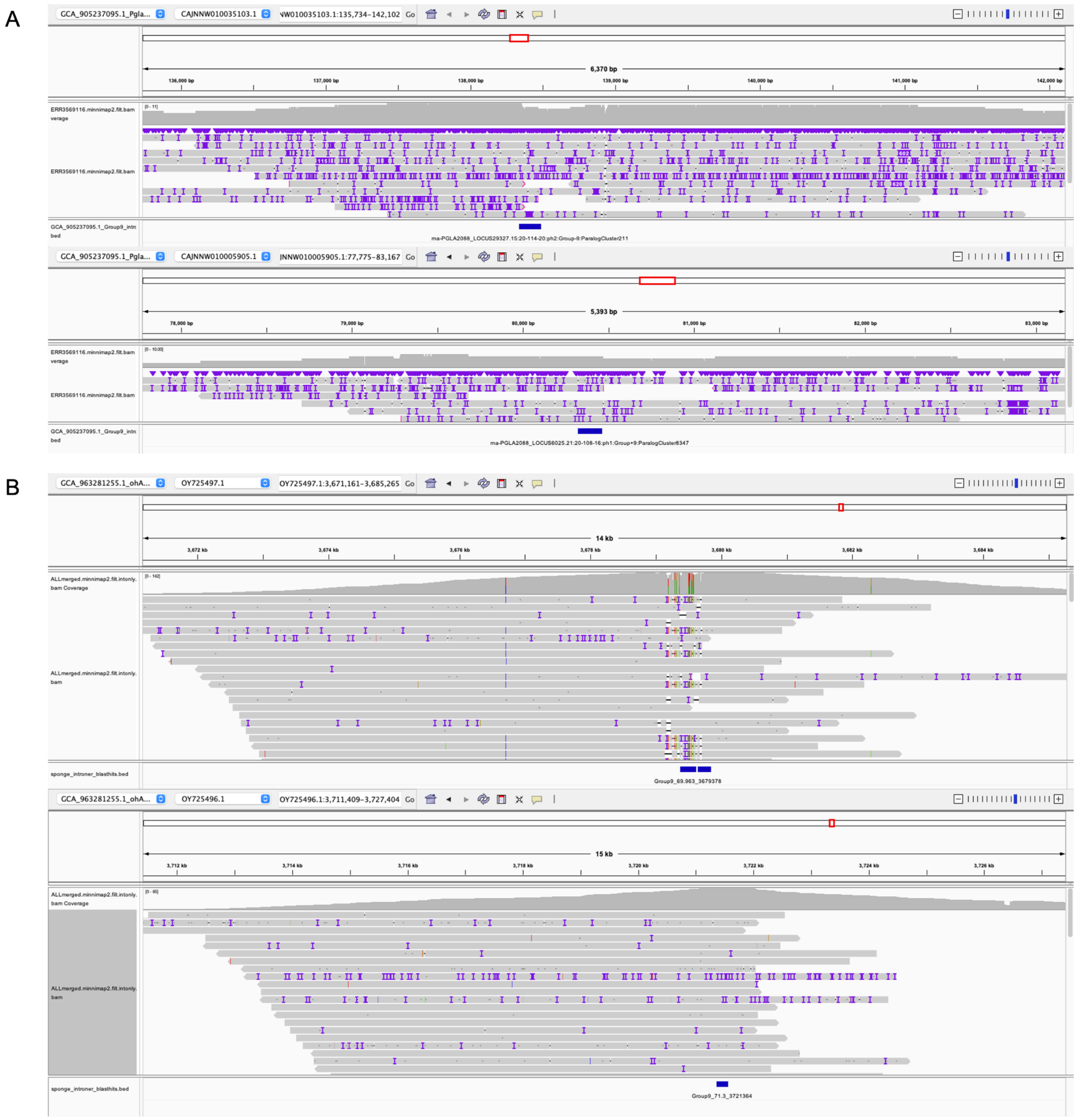


**fig. S12:** Verification of HGT between *Polarella glacialis* (GCA_905237095.1, introner Fam9) and *Aphrocallistes beatrix* (glass sponge). **(A)** Two examples of introner locations in *P. glacialis* are shown in blue, along with mapped long reads (PacBio), many of which span the introner region. **(B)** Two examples of introner blast hit locations in *A. beatrix* are shown in blue, along with mapped long reads (PacBio), many of which span the putative transferred region. Only reads overlapping the introner or blast hit regions of interest are shown.

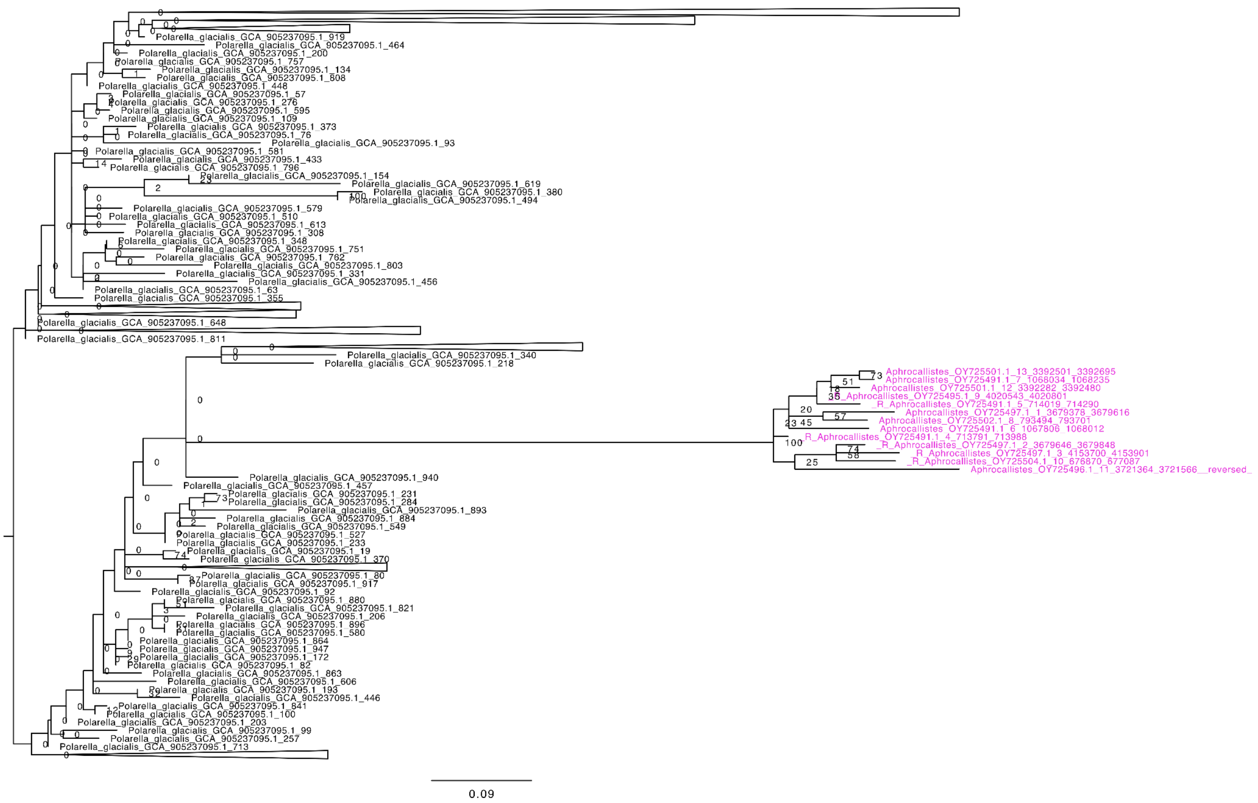

**fig. S13:** Maximum likelihood phylogeny for the introner element found in the dinoflagellate *Polarella* and the sponge *Aphrocallistes*, rooted at the midpoint. Bootstrap values out of 100 are indicated at the nodes. Scale bar measures substitutions per site.

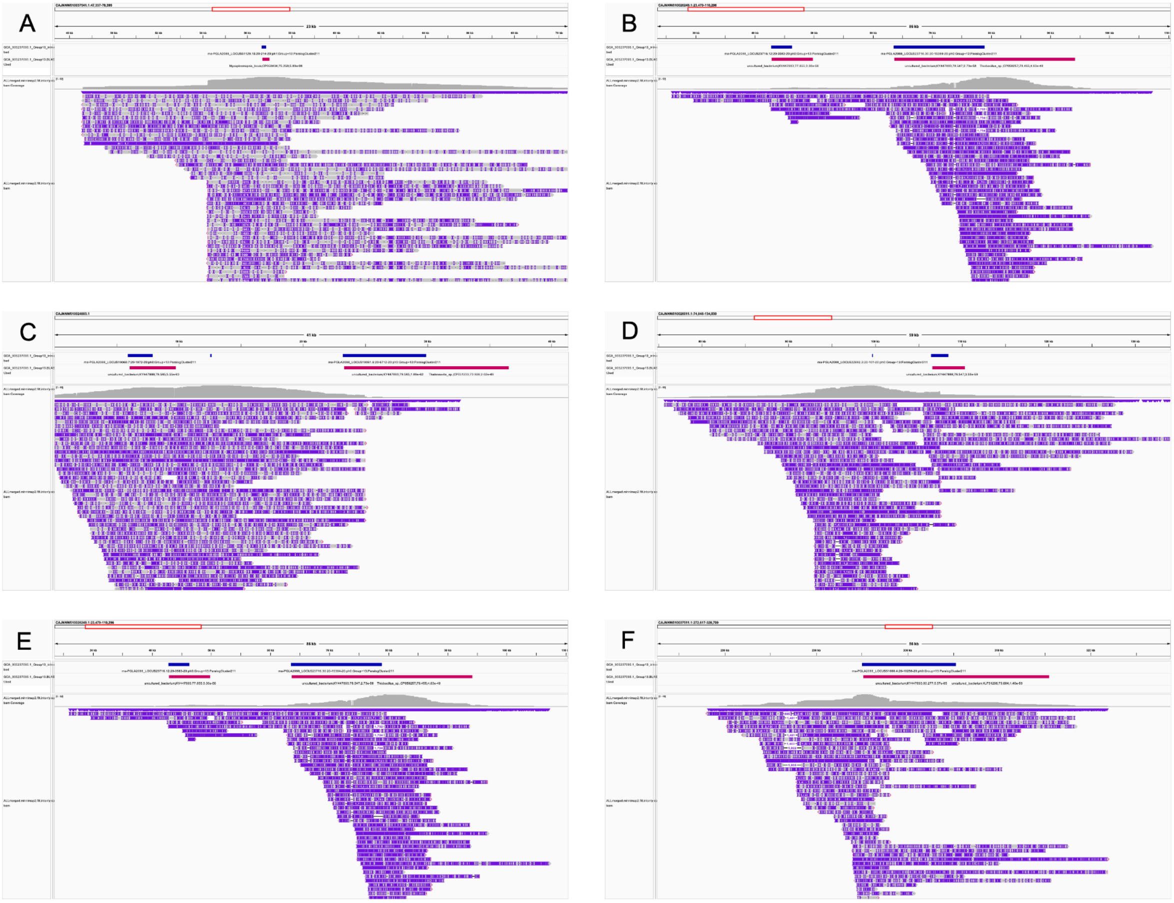

**fig. S14:** Verification of bacterial HGT to *Polarella glacialis* strain CCMP2088 (GCA_905237095.1, Fam13). Introner and bacterial blast hits are shown in blue and red respectively, along with mapped long reads (PacBio), many of which span the introner region. Only reads overlapping the introner are shown.

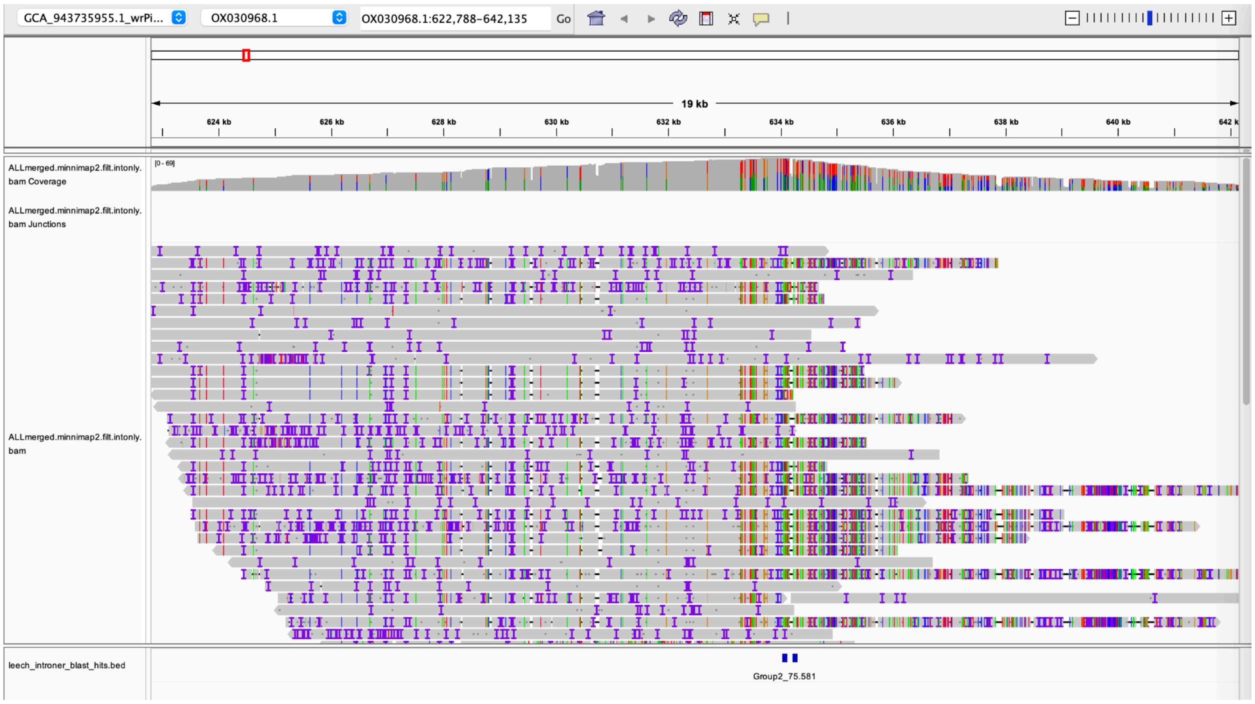


**fig. S15:** Verification of HGT between *Thalassiosira oceanica* (GCA_000296195.2) and *Piscicola geometra* (GCA_000296195.2). Locations of non-intron-generating homologs of *T. oceanica* introners in *P. geometra* are shown in blue (bottom track), along with mapped long reads (PacBio), many of which span the introner region.

#### Supplemental Tables

**table S1:** Species, NCBI accession and number of introner families for each surveyed genome.

**table S2:** Metadata, classification, and supporting information for each introner family identified. Columns A-E display species, phylogroup (common name), NCBI genome accession, family name and number of introners for each identified introner family. Column F shows RepeatClassifier TE classification. Column G shows the best hit for translated blasts against all TE proteins in the RepBase and DFAM databases. Column H shows all features identified by interproscan across all introner copies for each family as well as the number of copies in which that feature was identified (separated by a "_"). For example, "DUF3671_5" means that DUF3671 was present in 5 copies for a given introner family. Column I shows all features identified in each introner consensus sequence by hmmscans using all models available in PFAM. Column J shows the same but for all models available in the Gypsy database. Column K-M show MCHelper classifications and feature annotations for each introner consensus. Column K shows MCHelper results using the full introner consensus as input. Column L shows MCHelper results when only half of the introner consensus was used as input, and Column M shows results when only the first 100 bp were used. Columns N-O show conserved TE termini (retrieved from DFAM) identified using the original introner consensus sequence and the extended consensus produced by MCHelper, respectively. Columns P-Q display final classifications for each family after manual inspection. Column R shows additional notes about introner sequences. Columns S-U show information about the number and proportion of introners in the most common paralog family. Columns V-Y show information about introner splice sites.

**table S3:** Supporting information for Fig. 2C-E. Columns A-G show scaffold, position, introner family, intron number relative to transcription start site, species, locus tag and assembly accession, respectively, for each introner displayed in Fig. 2C-E.

**table S4:** Blast results for introner HGT candidates.

**table S5:** Summary of all identified introner HGT events. Columns A and B show species between which we observe HGT of introner sequences. Species which contain active introners are highlighted in green (We observe some cases in which introners are only actively generating introns in one lineage, and presumably functioning as ordinary TEs in the other). Columns C and D show TimeTree divergence times between each lineage displaying HGT in millions of years and supporting citations (*29*).

**table S6:** Taxa in Ascomycota TimeTree from Figure 3A in which introners are present, absent, or for which we have no data. The species (and subspecies) present/absent (column B) and species absent (column D) counts were tallied from unique species annotations for each genus in table S1.

**table S7:** Taxa in Eukaryota TimeTree from Figure 3D in which introners are present, absent, or for which we have no data. The species (and subspecies) present/absent (column B) and species absent (column D) counts were tallied from unique species annotations for each genus in table S1.

**table S8:** For each introner in *Polarella glacialis* strain CCMP2088 Fam9, the number of reads from each SRA accession that map spanning the full introner region.

**table S9:** Verification of horizontal transfer of *Polarella glacialis* Fam9 introner sequences between *Polarella glacialis* and *Aphrocallistes beatrix* (sponge). The number of reads from the available SRA accession that map spanning each *Polarella* introner blast hit region in *A. beatrix* are shown.

**table S10:** tblastn results for introner-containing genes against all viruses in RVDB (*90*). Only hits with e-value < 0.00001 are shown. In cases where a gene had multiple hits that passed this threshold, the hit with the lowest e-value is shown.

**table S11:** For each introner in *Strongylocentrotus purpuratus* Fam1, the number of reads from each SRA accession that map spanning the full introner region.

**table S12:** Verification of horizontal transfer of *Thalassiosira oceanica* Fam2 introner sequences between *T. oceanica* and *Piscicola geometra* (leach). The number of reads from the available SRA accession that map spanning each *T. oceanica* introner blast hit region in *P. geometra* are shown.

**table S13:** For each introner in *Polarella glacialis* strain CCMP2088 Fam13, the number of reads from each SRA accession that map spanning the full introner region, for confirmation of bacterial insertions.

#### References


57. C. Camacho, G. Coulouris, V. Avagyan, N. Ma, J. Papadopoulos, K. Bealer, T. L. Madden, BLAST+: architecture and applications. *BMC Bioinformatics* **10**, 421 (2009).

58. B. Buchfink, C. Xie, D. H. Huson, Fast and sensitive protein alignment using DIAMOND. *Nat. Methods* **12**, 59–60 (2015).

59. J. M. Storer, R. Hubley, J. Rosen, A. F. A. Smit, Curation guidelines for de novo generated transposable element families. *Curr Protoc* **1**, e154 (2021).

60. K. Katoh, D. M. Standley, MAFFT Multiple sequence alignment software version 7: improvements in performance and usability. *Mol. Biol. Evol.* **30**, 772–780 (2013).

61. A. Larsson, AliView: a fast and lightweight alignment viewer and editor for large datasets. *Bioinformatics* **30**, 3276–3278 (2014).

62. M. Burset, I. A. Seledtsov, V. V. Solovyev, Analysis of canonical and non-canonical splice sites in mammalian genomes. *Nucleic Acids Res.* **28**, 4364–4375 (2000).

63. B. Pucker, S. F. Brockington, Genome-wide analyses supported by RNA-Seq reveal non-canonical splice sites in plant genomes. *BMC Genomics* **19**, 980 (2018).

64. J. M. Flynn, R. Hubley, C. Goubert, J. Rosen, A. G. Clark, C. Feschotte, A. F. Smit, RepeatModeler2 for automated genomic discovery of transposable element families. *Proc. Natl. Acad. Sci. U. S. A.* **117**, 9451–9457 (2020).

65. R. Hubley, R. D. Finn, J. Clements, S. R. Eddy, T. A. Jones, W. Bao, A. F. A. Smit, T. J. Wheeler, The Dfam database of repetitive DNA families. *Nucleic Acids Res.* **44**, D81–9 (2016).

66. J. Jurka, Repbase update: a database and an electronic journal of repetitive elements. *Trends Genet.* **16**, 418–420 (2000).

67. C. Goubert, R. J. Craig, A. F. Bilat, V. Peona, A. A. Vogan, A. V. Protasio, A beginner's guide to manual curation of transposable elements. *Mob. DNA* **13**, 7 (2022).

68. B. J. Woodcroft, J. A. Boyd, G. W. Tyson, OrfM: a fast open reading frame predictor for metagenomic data. *Bioinformatics* **32**, 2702–2703 (2016).

69. P. Jones, D. Binns, H.-Y. Chang, M. Fraser, W. Li, C. McAnulla, H. McWilliam, J. Maslen, A. Mitchell, G. Nuka, S. Pesseat, A. F. Quinn, A. Sangrador-Vegas, M. Scheremetjew, S.-Y. Yong, R. Lopez, S. Hunter, InterProScan 5: genome-scale protein function classification. *Bioinformatics* **30**, 1236–1240 (2014).

70. M. Blum, H.-Y. Chang, S. Chuguransky, T. Grego, S. Kandasaamy, A. Mitchell, G. Nuka, T. Paysan-Lafosse, M. Qureshi, S. Raj, L. Richardson, G. A. Salazar, L. Williams, P. Bork, A. Bridge, J. Gough, D. H. Haft, I. Letunic, A. Marchler-Bauer, H. Mi, D. A. Natale, M. Necci, C. A. Orengo, A. P. Pandurangan, C. Rivoire, C. J. A. Sigrist, I. Sillitoe, N. Thanki, P. D. Thomas, S. C. E. Tosatto, C. H. Wu, A. Bateman, R. D. Finn, The InterPro protein families and domains database: 20 years on. *Nucleic Acids Res.* **49**, D344–D354 (2021).

71. S. R. Eddy, Accelerated profile HMM searches. *PLoS Comput. Biol.* **7**, e1002195 (2011).

72. J. Mistry, S. Chuguransky, L. Williams, M. Qureshi, G. A. Salazar, E. L. L. Sonnhammer, S. C. E. Tosatto, L. Paladin, S. Raj, L. J. Richardson, R. D. Finn, A. Bateman, Pfam: The protein families database in 2021. *Nucleic Acids Res.* **49**, D412–D419 (2021).

73. C. Llorens, R. Futami, L. Covelli, L. Domínguez-Escribá, J. M. Viu, D. Tamarit, J. Aguilar-Rodríguez, M. Vicente-Ripolles, G. Fuster, G. P. Bernet, F. Maumus, A. Munoz-Pomer, J. M. Sempere, A. Latorre, A. Moya, The Gypsy Database (GyDB) of mobile genetic elements: release 2.0. *Nucleic Acids Res.* **39**, D70–4 (2011).

74. S. Orozco-Arias, P. Sierra, R. Durbin, J. González, MCHelper automatically curates transposable element libraries across species, *bioRxiv* (2023)p. 2023.10.17.562682.

75. T. J. Wheeler, S. R. Eddy, nhmmer: DNA homology search with profile HMMs. *Bioinformatics* **29**, 2487–2489 (2013).

76. D. M. Goodstein, S. Shu, R. Howson, R. Neupane, R. D. Hayes, J. Fazo, T. Mitros, W. Dirks, U. Hellsten, N. Putnam, D. S. Rokhsar, Phytozome: a comparative platform for green plant genomics. *Nucleic Acids Res.* **40**, D1178–86 (2012).

77. N. A. O'Leary, M. W. Wright, J. R. Brister, S. Ciufo, D. Haddad, R. McVeigh, B. Rajput, B. Robbertse, B. Smith-White, D. Ako-Adjei, A. Astashyn, A. Badretdin, Y. Bao, O. Blinkova, V. Brover, V. Chetvernin, J. Choi, E. Cox, O. Ermolaeva, C. M. Farrell, T. Goldfarb, T. Gupta, D. Haft, E. Hatcher, W. Hlavina, V. S. Joardar, V. K. Kodali, W. Li, D. Maglott, P. Masterson, K. M. McGarvey, M. R. Murphy, K. O'Neill, S. Pujar, S. H. Rangwala, D. Rausch, L. D. Riddick, C. Schoch, A. Shkeda, S. S. Storz, H. Sun, F. Thibaud-Nissen, I. Tolstoy, R. E. Tully, A. R. Vatsan, C. Wallin, D. Webb, W. Wu, M. J. Landrum, A. Kimchi, T. Tatusova, M. DiCuccio, P. Kitts, T. D. Murphy, K. D. Pruitt, Reference sequence (RefSeq) database at NCBI: current status, taxonomic expansion, and functional annotation. *Nucleic Acids Res.* **44**, D733–45 (2016).

78. Y. Xiong, T. H. Eickbush, Origin and evolution of retroelements based upon their reverse transcriptase sequences. *EMBO J.* **9**, 3353–3362 (1990).

79. P. J. A. Cock, T. Antao, J. T. Chang, B. A. Chapman, C. J. Cox, A. Dalke, I. Friedberg, T. Hamelryck, F. Kauff, B. Wilczynski, M. J. L. de Hoon, Biopython: freely available Python tools for computational molecular biology and bioinformatics. *Bioinformatics* **25**, 1422–1423 (2009).

80. J. Trifinopoulos, L.-T. Nguyen, A. von Haeseler, B. Q. Minh, W-IQ-TREE: a fast online phylogenetic tool for maximum likelihood analysis. *Nucleic Acids Res.* **44**, W232–5 (2016).

81. J. L. Steenwyk, T. J. Buida 3rd, Y. Li, X.-X. Shen, A. Rokas, ClipKIT: A multiple sequence alignment trimming software for accurate phylogenomic inference. *PLoS Biol.* **18**, e3001007 (2020).

82. B. Q. Minh, H. A. Schmidt, O. Chernomor, D. Schrempf, M. D. Woodhams, A. von Haeseler, R. Lanfear, IQ-TREE 2: New models and efficient methods for phylogenetic inference in the genomic era. *Mol. Biol. Evol.* **37**, 1530–1534 (2020).

83. P. Sagulenko, V. Puller, R. A. Neher, TreeTime: Maximum-likelihood phylodynamic analysis. *Virus Evol* **4**, vex042 (2018).

84. T. Kasuga, T. J. White, J. W. Taylor, Estimation of nucleotide substitution rates in Eurotiomycete fungi. *Mol. Biol. Evol.* **19**, 2318–2324 (2002).

85. M. L. Berbee, J. W. Taylor, Dating the molecular clock in fungi – how close are we? *Fungal Biol. Rev.* **24**, 1–16 (2010).

86. J. Labbé, C. Murat, E. Morin, G. A. Tuskan, F. Le Tacon, F. Martin, Characterization of transposable elements in the ectomycorrhizal fungus Laccaria bicolor. *PLoS One* **7**, e40197

(2012).

87. K. Silliman, J. L. Indorf, N. Knowlton, W. E. Browne, C. Hurt, Base-substitution mutation rate across the nuclear genome of Alpheus snapping shrimp and the timing of isolation by the Isthmus of Panama. *BMC Ecol Evol* **21**, 104 (2021).

88. R. Allio, S. Donega, N. Galtier, B. Nabholz, Large variation in the ratio of mitochondrial to nuclear mutation rate across animals: implications for genetic diversity and the use of mitochondrial DNA as a molecular marker. *Mol. Biol. Evol.* **34**, 2762–2772 (2017).

89. M. Krasovec, S. Sanchez-Brosseau, G. Piganeau, First estimation of the spontaneous mutation rate in diatom. *Genome Biol. Evol.* **11**, 1829–1837 (2019).

90. N. Goodacre, A. Aljanahi, S. Nandakumar, M. Mikailov, A. S. Khan, A reference viral database (RVDB) to enhance bioinformatics analysis of high-throughput sequencing for novel virus detection. *mSphere* **3** (2018).

91. C. L. Schoch, S. Ciufo, M. Domrachev, C. L. Hotton, S. Kannan, R. Khovanskaya, D. Leipe, R. Mcveigh, K. O'Neill, B. Robbertse, S. Sharma, V. Soussov, J. P. Sullivan, L. Sun, S. Turner, I. Karsch-Mizrachi, NCBI Taxonomy: a comprehensive update on curation, resources and tools. *Database* **2020** (2020).

92. J. Huerta-Cepas, F. Serra, P. Bork, ETE 3: Reconstruction, analysis, and visualization of phylogenomic data. *Mol. Biol. Evol.* **33**, 1635–1638 (2016).

93. I. Letunic, P. Bork, Interactive Tree of Life (iTOL) v6: recent updates to the phylogenetic tree display and annotation tool. *Nucleic Acids Res.*, doi: 10.1093/nar/gkae268 (2024).

94. D. A. R. Eaton, Toytree: A minimalist tree visualization and manipulation library for Python. *Methods Ecol. Evol.* **11**, 187–191 (2020).

95. H. Li, Minimap2: pairwise alignment for nucleotide sequences. *Bioinformatics* **34**, 3094–3100 (2018).

96. A. R. Quinlan, I. M. Hall, BEDTools: a flexible suite of utilities for comparing genomic features. *Bioinformatics* **26**, 841–842 (2010).

97. J. T. Robinson, H. Thorvaldsdóttir, W. Winckler, M. Guttman, E. S. Lander, G. Getz, J. P. Mesirov, Integrative genomics viewer. *Nat. Biotechnol.* **29**, 24–26 (2011).

98. M. M. Teixeira, L. F. Moreno, B. J. Stielow, A. Muszewska, M. Hainaut, L. Gonzaga, A. Abouelleil, J. S. L. Patané, M. Priest, R. Souza, S. Young, K. S. Ferreira, Q. Zeng, M. M. L. da Cunha, A. Gladki, B. Barker, V. A. Vicente, E. M. de Souza, S. Almeida, B. Henrissat, A. T. R. Vasconcelos, S. Deng, H. Voglmayr, T. A. A. Moussa, A. Gorbushina, M. S. S. Felipe, C. A. Cuomo, G. S. de Hoog, Exploring the genomic diversity of black yeasts and relatives (Chaetothyriales, Ascomycota). *Stud. Mycol.* **86**, 1–28 (2017).

99. X.-X. Shen, J. L. Steenwyk, A. L. LaBella, D. A. Opulente, X. Zhou, J. Kominek, Y. Li, M.

Groenewald, C. T. Hittinger, A. Rokas, Genome-scale phylogeny and contrasting modes of genome evolution in the fungal phylum Ascomycota. *Sci Adv* **6** (2020).

100. E. Kraichak, J.-P. Huang, M. Nelsen, S. D. Leavitt, H. T. Lumbsch, A revised classification of orders and families in the two major subclasses of Lecanoromycetes (Ascomycota) based on a temporal approach. *Bot. J. Linn. Soc.* **188**, 233–249 (2018).

101. P. K. Divakar, A. Crespo, M. Wedin, S. D. Leavitt, D. L. Hawksworth, L. Myllys, B. McCune, T. Randlane, J. W. Bjerke, Y. Ohmura, I. Schmitt, C. G. Boluda, D. Alors, B. Roca-Valiente, R. Del-Prado, C. Ruibal, K. Buaruang, J. Núñez-Zapata, G. Amo de Paz, V. J. Rico, M. C. Molina, J. A. Elix, T. L. Esslinger, I. K. K. Tronstad, H. Lindgren, D. Ertz, C. Gueidan, L. Saag, K. Mark, G. Singh, F. Dal Grande, S. Parnmen, A. Beck, M. N. Benatti, D. Blanchon, M. Candan, P. Clerc, T. Goward, M. Grube, B. P. Hodkinson, J.-S. Hur, G. Kantvilas, P. M. Kirika, J. Lendemer, J.-E. Mattsson, M. I. Messuti, J. Miadlikowska, M. Nelsen, J. I. Ohlson, S. Pérez-Ortega, A. Saag, H. J. M. Sipman, M. Sohrabi, A. Thell, G. Thor, C. Truong, R. Yahr, D. K. Upreti, P. Cubas, H. T. Lumbsch, Evolution of complex symbiotic relationships in a morphologically derived family of lichen-forming fungi. *New Phytol.* **208**, 1217–1226 (2015).
